# SMCHD1 is a novel target for gene-activation therapy to treat Prader-Willi Syndrome

**DOI:** 10.64898/2026.05.13.725051

**Authors:** Megan Iminitoff, Anna Le Fevre, Tamara Cameron, Hannah K Vanyai, Caleb Chew, Sarah A Kinkel, Kelsey Breslin, Susanne Theiß, Quentin Gouil, Merlin Thomas, James M Murphy, Christian P. Schaaf, Andrew Keniry, Marnie E Blewitt

## Abstract

Prader-Willi Syndrome (PWS) is a neurodevelopmental disorder caused by lack of gene expression from the active paternal allele at an imprinted gene cluster on chromosome 15. Current treatments have limited efficacy as they target individual symptoms rather than the underlying cause of disease. All patients preserve a normal, yet epigenetically-silenced, copy of the PWS cluster genes; activation of this imprinted copy to restore necessary gene expression is an appealing option for tackling the root of the disorder. Here we have addressed the potential to activate these silent maternal genes by targeting the epigenetic regulator Structural Maintenance of Chromosomes Hinge domain containing 1 (SMCHD1). First, we expanded the role of SMCHD1 in repressing the PWS cluster from mice to humans, a critical step if SMCHD1 is to be a drug target. Second, we discovered that SMCHD1 represses the entire PWS locus in neural lineages, extending its previously known role at only half of the PWS genes. We show that deleting *Smchd1* after early development *in vivo* is effective at causing PWS gene-activation in disease-relevant mouse tissues including hypothalamus, and that this has beneficial effects on phenotypes observed in a PWS mouse model. Despite SMCHD1 having a role in gene silencing elsewhere in the genome, our data suggest that targeting SMCHD1 after early development is remarkably safe. Taken together, these data propose SMCHD1 as a novel target for gene-activation therapy for PWS.

## INTRODUCTION

Prader-Willi Syndrome (PWS) is a neurodevelopmental disorder with a prevalence of 1:10,000 - 1:30,000 live births^1^. Infants initially present with hypotonia and feeding difficulties, and early molecular diagnosis usually occurs within the first 2 months of life, although this varies by location and access to resources^2,3^. Patients living with PWS develop a variety of metabolic, endocrine, cognitive, and behavioural problems including intellectual disability, sex and growth hormone deficiencies, hyperphagia, and obesity. Additional variable features include scoliosis, sleep and respiratory issues and psychiatric disorders^4–6^. Current best practise in the management of PWS is predominantly symptomatic and supportive, targeting individual clinical manifestations rather than the underlying genetic aetiology. Consequently, no single intervention provides comprehensive disease modification; treatments must be tailored to individual patients and are frequently subject to contraindications^10^.

PWS symptoms stem from a lack of gene expression at the imprinted 15q11-q13 locus. This locus contains a cluster of maternally imprinted genes, which are exclusively paternally expressed in the brain, and one paternally imprinted gene (*UBE3A*), which is exclusively maternally expressed in the brain^11^. PWS arises when the paternally expressed PWS cluster genes are not inherited due to either deletions (∼60% of patients) or maternal uniparental disomy (mUPD) (∼36% of patients). The remaining ∼4% of patients have a present but epigenetically silenced paternal allele due to atypical deletions or defects in the primary imprint^8,12^. Angelman Syndrome is the reciprocal neurodevelopmental imprinting disorder, that arises if expression of the maternal copy of the gene *UBE3A* is disrupted^13,14^.

The PWS cluster hosts a DNA methylation primary imprint at its imprinting control region (ICR) that controls the imprinted expression of the whole gene cluster. This ICR is bipartite, with the PWS ICR methylated on the maternal allele and unmethylated on the paternal allele. The adjacent Angelman Syndrome ICR is thought to suppress the activity of the PWS ICR in the maternal germline and is subordinate in most cells^15,16^. Another critical mechanism of imprinting control at the 15q11-q13 region is the long non-coding RNA (lncRNA) Small Nucleolar Host Gene 14 (SNHG14). This lncRNA originates from a shared upstream promoter of *SNURF-SNRPN* and serves as the host gene for multiple transcripts, not all of which are yet annotated, including the small nucleolar RNAs (snoRNAs) and the antisense transcript for *UBE3A* (*UBE3A-ATS*)^17^. The proximal portion of *SNHG14* is expressed from the active paternal allele in all tissues; however, the distal end containing *UBE3A-ATS* is expressed only in neurons. Transcription of *SNHG14* through the snoRNAs, their host genes, and *UBE3A-ATS* results in interruption of the *UBE3A* transcript in the opposite direction and is widely accepted to be the mechanism of imprinting control for *UBE3A* in neurons^18–20^. The *UBE3A* promoter itself is unmethylated on both parental alleles, allowing for biallelic expression in other tissues where transcription of the distal end of *SNHG14* does not occur^21^. Both the genes and the known mechanisms of imprinting control at the PWS cluster are conserved with remarkable synteny on chromosome 7 in mice^22^.

Given that gene expression is controlled through epigenetic processes at the PWS locus, and that all PWS patients have the maternal copy of PWS locus genes, epigenetic-based methods of gene-activation are an attractive therapeutic intervention for PWS. Unlike current therapeutics, gene-activation therapy would target the root cause of the disorder and therefore offer hope of a disease-modifying therapy. Investigation into such therapies has explored inhibition of DNA methyltransferases to demethylate the locus for gene-activation^23^. More recent studies have uncovered additional repressive factors acting on the maternal allele and look at targeting these factors to trigger maternal gene-activation^15,24–26^. Currently there are no epigenetic therapies in clinical trial for PWS, in part due to the factors in question having critical roles throughout the genome and therefore creating toxicity issues^30^. By contrast, clinical trials have been launched in recent years for antisense oligonucleotides (ASOs) targeting *UBE3A-ATS* for the treatment of Angelman Syndrome^31–34^ (ClinicalTrials.gov, NCT06614606, NCT06617426, NCT06415344, NCT07136454, NCT04259281, NCT04428281).

The focus of gene-activation studies for PWS has been on the distal PWS cluster genes *SNURF-SNRPN* and the *SNORD* snoRNAs^15,24–28^. This may be due to reports of patients with small, atypical deletions involving parts of the SNHG14 non-coding RNA but not the PWS ICR, which suggested a possible ‘minimal critical region’ for PWS surrounding the SNORD116 snoRNAs^35–41^. Though the SNORD116 snoRNAs and their targets are likely involved in PWS pathogenesis, individuals with the smallest deletions^36,40^, appear atypical and clinically relatively mild for PWS, such as achieving normal adult height without growth hormone support. These reports indicate that deficiency of the SNORD116 snoRNAs alone may not result in a typical PWS phenotype. Further, there is a related disorder, Schaaf-Yang Syndrome, caused by truncating variants in MAGEL2 where loss of paternal *MAGEL2* and the presence of a truncated protein result in overlapping features with PWS^42–45^. *MKRN3*, another gene in the proximal region of the cluster, is associated with central precocious puberty^46^. These disorders suggest that lack of expression of proximal PWS cluster genes likely also contributes to the PWS phenotype.

In this study we investigated Structural Maintenance of Chromosomes Hinge Domain Containing 1 (SMCHD1) as a potential target for gene-activation therapy for PWS. SMCHD1 is a chromosomal ATPase consisting of a C-terminal SMC hinge domain, and an N-terminal GHKL-type ATPase, which enable protein dimerisation and interaction with chromatin crucial to its function as an epigenetic repressor^47–54^. SMCHD1 is considered a non-canonical member of the structural maintenance of chromosomes (SMC) family of proteins due to its distinct architecture. SMCHD1 plays an essential role during early development where it is required for silencing the inactive X chromosome in females and several other autosomal clustered gene families, such as the *Hox* gene clusters important in embryo patterning, clustered protocadherins and select imprinted regions^55–61^.

We and others have shown SMCHD1 suppresses the maternal copy of proximal PWS genes *Magel2*, *Ndn* and *Mkrn3* in mice^51,58–62^. Interestingly, absence of SMCHD1 does not alter the primary DNA methylation imprint, even when *Smchd1* deletion occurs in the female germline^60^, suggesting SMCHD1 could play a role downstream of the methylation imprint, and therefore after early development. Furthermore, deletion of *Smchd1* from mid-gestation in the blood *in vivo*, resulted in upregulation of *Mkrn3* and *Peg12* (a mouse-specific PWS locus gene) in B cells^61^, while deletion of *Smchd1* in mid-gestation cortical neural stem cells *in vitro* resulted in activation of *Ndn*^58^. These data suggest it may be effective to target SMCHD1 after early development. In both cases, while PWS locus genes were activated, there was no change in expression from the inactive X chromosome. Together these data give an indication that targeting SMCHD1 after early development could be effective in gene-activation without disrupting developmental processes such as X-chromosome inactivation and *Hox* gene silencing. Additionally, the fact that SMCHD1 is an enzyme^46^ makes it a potentially viable therapeutic target.

In this study we sought to further investigate the plausibility of targeting SMCHD1 to treat PWS. We showed that SMCHD1 represses human maternal PWS gene expression in a PWS patient-derived neural system, and expanded SMCHD1’s role to silencing of the distal cluster genes. Through generation of *in vivo* mouse models, we addressed potential safety and functional consequences of targeting SMCHD1 for gene-activation therapy after its critical role in X-inactivation and *Hox* gene silencing. We found minimal genome-wide effects of targeting SMCHD1 at this developmental time point and no concerning functional effects in our behavioural cohorts. Furthermore, behavioural tests suggest functional improvements to PWS phenotypes upon *Smchd1* deletion in the brain. Taken together these data provide a robust foundation for continued pursuit of SMCHD1 as a target for gene-activation therapy to treat PWS.

## RESULTS

### SMCHD1 is required for maintenance of imprinting at the PWS locus in humans

To begin investigating whether the known role of SMCHD1 in maintenance of imprinting at the PWS cluster in mice^51,58–62^ translates to human (**Fig. 1A**) we created clonal *SMCHD1* knockout HEK263T cell lines using CRISPR-*Cas9* editing (**Fig. 1B, S1A, Supplementary Table 1**). By RT-qPCR we found that in *SMCHD1* knockout cells there was a significant increase in *MAGEL2* expression compared with *SMCHD1* wild-type clonal lines. We found no change in expression of the distal PWS cluster gene *SNRPN* (**Fig. 1C**) in line with our prior data in mouse^51,58–62^ suggesting that at least in these contexts SMCHD1’s major role may be at the proximal end of the PWS cluster.

**Figure 1:**
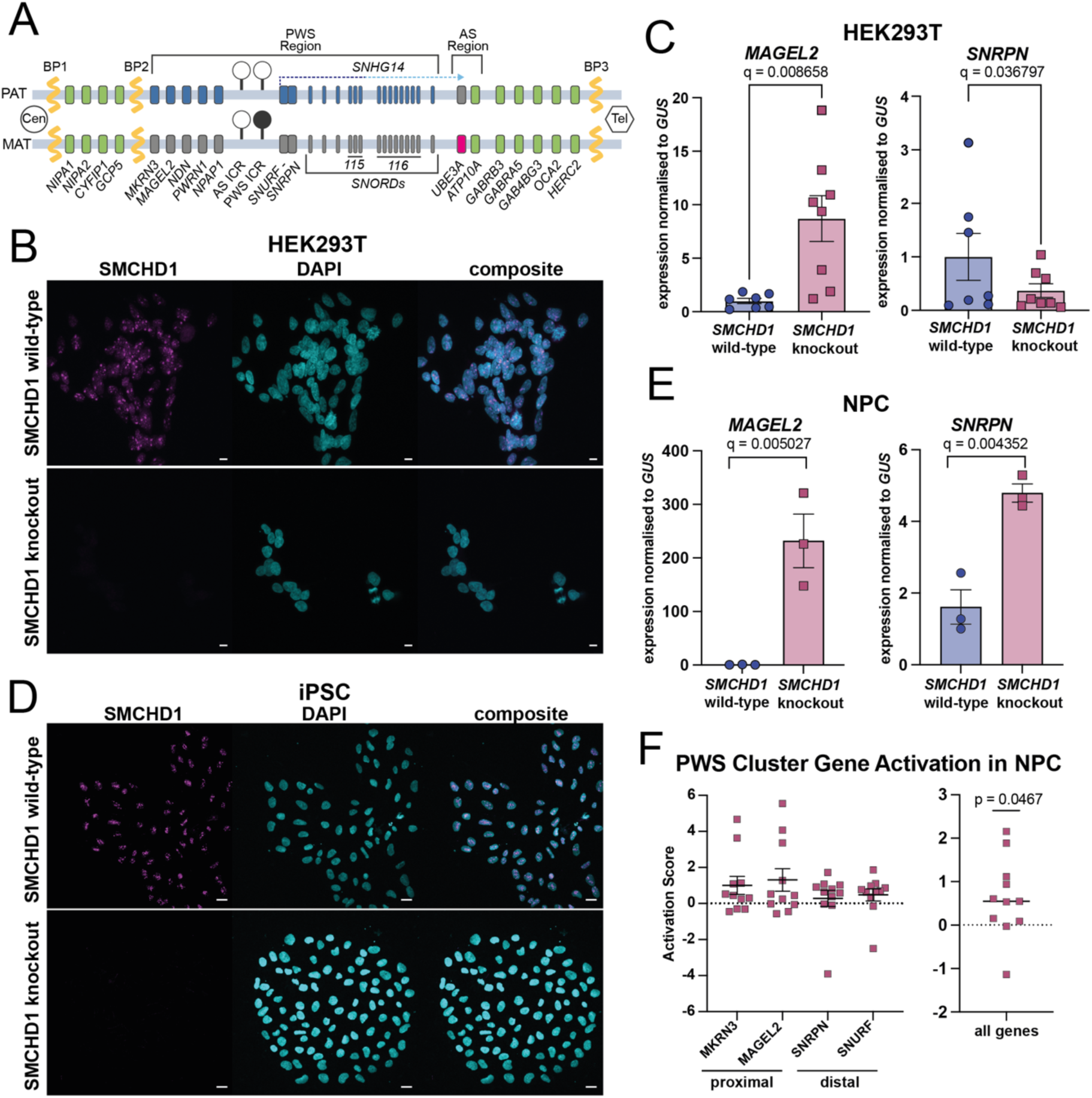
SMCHD1 deletion leads to maternal PWS gene-activation in human neural cell models of PWS. **A)** Schematic representation of the human 15q11-q13 region, common breakpoints (BP) found in patients with PWS are represented by a yellow zig-zag, biallelically expressed non-imprinted genes in green, maternally expressed imprinted genes in pink, paternally expressed imprinted genes in blue, repressed copy of imprinted genes in grey. Note this is reflective of expression status in neurons. Arrow depicts direction of transcription for lncRNA SNHG14, the darker blue portion of the arrow is transcribed in all tissues, the lighter blue portion is exclusively transcribed in neurons, methylated region represented by a filled circle, unmethylated by an open circle, Cen and Tel represent the direction of the centromere and telomere. **B/D)** Representative confocal images showing SMCHD1 protein by immunofluorescence in (**B**) HEK263T clonal lines and (**D**) mUPD iPSC, SMCHD1 in magenta, DAPI in cyan, scale bar 20*µ*m. **C)** PWS gene expression in clonal HEK263Ts by RT-qPCR, replicates from three clonal lines for each condition across multiple separate experiments, t-tests with Benjamini and Hochberg corrections for multiple testing, q < 0.01 for significance. **E)** PWS gene expression by RT-qPCR in NPCs differentiated from either SMCHD1 wild-type or SMCHD1 knockout PWS mUPD iPSCs, replicates from three separate clonal lines for each condition, t-tests with Benjamini and Hochberg corrections for multiple testing, q < 0.01 for significance. **F)** Graphs showing the average fold change by RNAseq for *MAGEL2*, *MKRN3*, *SNRPN* and *SNURF* in patient derived NPCs treated with an ASO to deplete SMCHD1 compared to a matched sample treated with a control ASO either separately for each gene (left panel) or as a single activation score for all genes (right panel). NPCs differentiated from both mUPD and LD PWS patient iPSCs, each data point is one replicate from either a unique batch of differentiated cells within a single experiment or from replicate experiments. One sample t-and Wilcoxon test for significance p < 0.05.

For a more powerful allele-specific analysis of the effect of *SMCHD1* deletion at the PWS cluster in humans we used the same guides to knock out *SMCHD1* in PWS patient-derived induced pluripotent stem cells (iPSCs) (**Fig. 1D, S1B, Supplementary Table 2**). Given these cells only possess imprinted maternal alleles, an increase in gene expression reflects activation of the typically silent maternal allele. Following differentiation into neural progenitor cells (NPCs) RT-qPCR revealed that both *MAGEL2* and *SNRPN* showed significantly increased expression in *SMCHD1* knockout cell lines compared with controls, albeit with a lower fold change for *SNRPN* (**Fig. 1E**). Activation of *SNRPN* as well as *MAGEL2* suggests that there may be a neural-specific role for SMCHD1 at the distal end of the PWS cluster.

To probe the developmental timing of SMCHD1 silencing at the locus, we knocked out *SMCHD1* in NPCs after differentiation from PWS patient-derived iPSCs (**Supplementary Table 2**). Given that *SMCHD1* knockout does not occur in every cell (**Fig. S1C**), we used immunofluorescent staining for SMCHD1 and SNRPN to assess SNRPN levels in the absence of SMCHD1 in NPCs. Available antibodies for the PWS cluster proteins are scarce; however, despite some background, using the chosen antibody and appropriate controls we observed a significant increase in SNRPN signal in the SMCHD1 positive nuclei compared with SMCHD1 negative controls (**Fig. S1D-E**). This reveals that targeting SMCHD1 at a later timepoint in development, once the neural lineage is already specified, is still effective at maternal PWS gene-activation.

Using the same patient-derived NPC model we took a complementary approach and assessed SMCHD1 depletion via treatment with vivo-morpholino antisense oligonucleotides (ASOs) to deplete SMCHD1 by exon skipping (**Fig. S1F-G**). Across replicates we achieved variable activation of PWS genes in SMCHD1 depleted NPCs compared with controls, from both deletion and mUPD patient subtypes. Here we only report on the four cluster genes (*MAGEL2*, *MKRN3*, *SNURF* and *SNRPN*) with detectable expression in NPCs by RNA sequencing. Similar to earlier data, we see slightly more activation at proximal genes than at distal genes. When we quantify the overall effect at the PWS region by summarising activation of proximal and distal genes together this reaches statistical significance, indicating a whole-locus impact resulting from SMCHD1 depletion (**Fig. 1F**).

Together, these results indicate SMCHD1 function at the PWS cluster in humans is akin to previously published findings in mice with the expansion of its role to regulate the distal end of the PWS cluster.

### CNS-specific deletion of Smchd1 in vivo restores maternal PWS gene expression in the brain

To date, studies in mouse have examined the role of SMCHD1 at the PWS cluster (**Fig. 2A**) *in vitro,* and in embryos, placentae, whole brain and blood cells *in vivo*^51,58–62^. To confirm that loss-of-imprinting at the PWS locus resulting from *Smchd1* deletion holds true in PWS-relevant tissues *in vivo*, we crossed *Nestin*-Cre^63^ and our *Smchd1* floxed allele^64^ strains to generate a brain-specific deletion of *Smchd1* from embryonic day (E)10.5. The timing of deletion is importantly after SMCHD1’s crucial role in epigenetic silencing during the establishment of X-chromosome inactivation, and silencing of *Hox* genes and other autosomal targets^51,58,61,64,65^. We crossed this new brain-specific deletion model with a published *Magel2*-*LacZ* knockin-knockout model^66^, which carries a *LacZ* knockin at the endogenous *Magel2* promoter that functions as a reporter for *Magel2* expression (**Fig. 2B**). To test the effect of neural *Smchd1* deletion after early embryonic development on maternal *Magel2* expression we created a cohort of mice carrying the *Magel2*-*LacZ* reporter gene on the typically silent maternal allele and screened for *Magel2* expression using X-gal staining on fixed brain cryosections at approximately postnatal day 25. In the *Magel2*^patKI/+^ positive control brains there was observable *Magel2* signal (**Fig. 2C**), which was not seen in *Magel2*^matKI/+^ brains replete for SMCHD1(**Fig. 2D**). We saw substantial maternal *Magel2* activation in the *Magel2*^matKI/+^ samples with *Smchd1* deleted, although it did not reach normal paternal levels of *Magel2* in the brain, likely because activation does not occur in all cells (**Fig. 2E**). To remove SMCHD1 at the most therapeutically relevant time *in vivo* we attempted postnatal deletion of *Smchd1* with tamoxifen-inducible CreERT2^67^ in the first 7 days after birth. The tamoxifen doses required to achieve even partial deletion of *Smchd1* in the brain proved prohibitively toxic to C57BL/6J neonates; however, from two surviving mice we observed some maternal *Magel2* activation in the context of only partial *Smchd1* deletion (**Fig. 2F, S2A)**, with both *Magel2*^matKI/+^ and *Magel2*^patKI/+^ controls appearing as expected (**S2B-D**). These data provide proof-of-concept that *in vivo* deletion of *Smchd1* in the brain can activate maternal *Magel2* gene expression.

**Figure 2:**
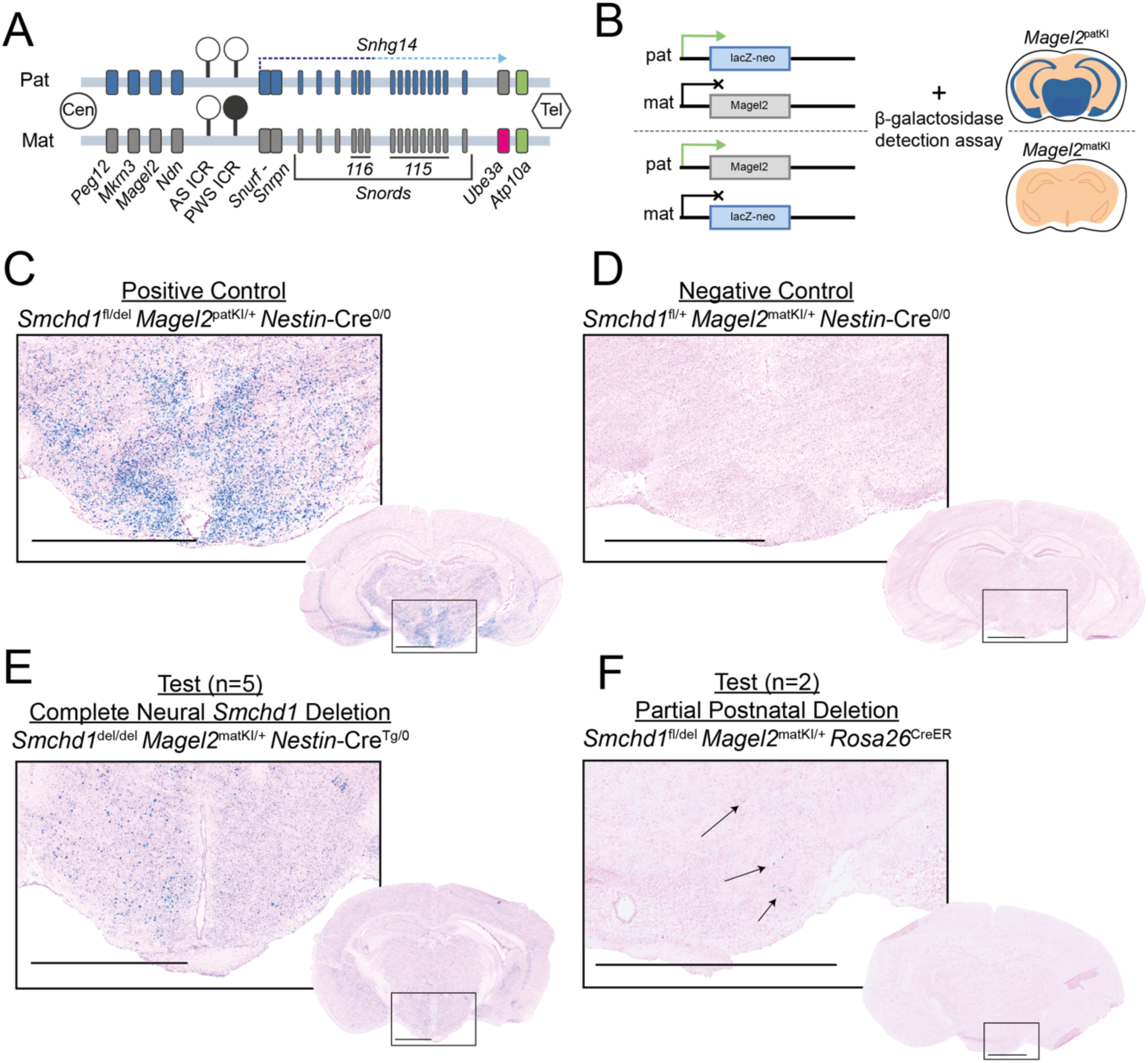
*Smchd1* deletion results in loss-of-imprinting and activation of maternal *Magel2* in mouse brain. **A)** Schematic of the PWS locus on mouse chromosome 7, paternally expressed genes in blue, maternally expressed genes in pink, biallelically expressed genes in green, repressed genes in grey, filled circle depicts methylated region, open circle represents unmethylated region. The arrow represents the direction of transcription for the lncRNA Snhg14 with dark blue representing the portion transcribed in all tissues and light blue representing the portion exclusively transcribed in neurons, Cen and Tel depict the direction of the centromere and telomere. **B)** Representation of *LacZ* reporter function in the *Magel2*-*LacZ* knockout-knockin mouse model when visualised by X-gal staining (blue). **C-F)** Representative images of brain cryosections stained with X-gal showing proxy for *Magel2* expression (blue) and counterstained with nuclear fast red (pink), zoomed images correspond to their overlayed whole brain slice, scale bars 1500 *µ*m. Note vertical brain sections are not matched exactly by slice. Test animals (**E**) n=5, (**F**) n=2.

To expand our understanding beyond *Magel2* we designed an RNA sequencing experiment to compare gene expression in the brain across the PWS cluster between *Smchd1*^del/del^ and *Smchd1*^fl/fl^ (representative of wild-type) mice in an allele-specific manner. C57BL/6J mice were intercrossed with Castaneus mice such that strain-specific SNPs would allow discernment between maternal and paternal alleles (**Fig. 3A**). We analysed the hypothalamus and cortex as PWS-relevant tissues in postnatal day 25 animals. We performed a genome-wide analysis of all reads, discovering only 18 significantly differentially expressed genes between the *Smchd1*^del/del^ and *Smchd1*^fl/fl^ hypothalamus tissue samples, one of these being *Smchd1* itself and only three being known *Smchd1* targets (*Pcdhb1*, *Pcdhb3*, *Pcdhb13*) (**Fig. 3B, Supplementary Table 3**). We observed 56 differentially expressed genes in the cortex (**Fig. 3C, Supplementary Table 4**). This relatively limited differential expression, even in the context of females where SMCHD1’s role in X inactivation means more changes may be expected, along with survival of healthy animals from both sexes, implies removal of *Smchd1* at this developmental stage is likely to be safe.

**Figure 3:**
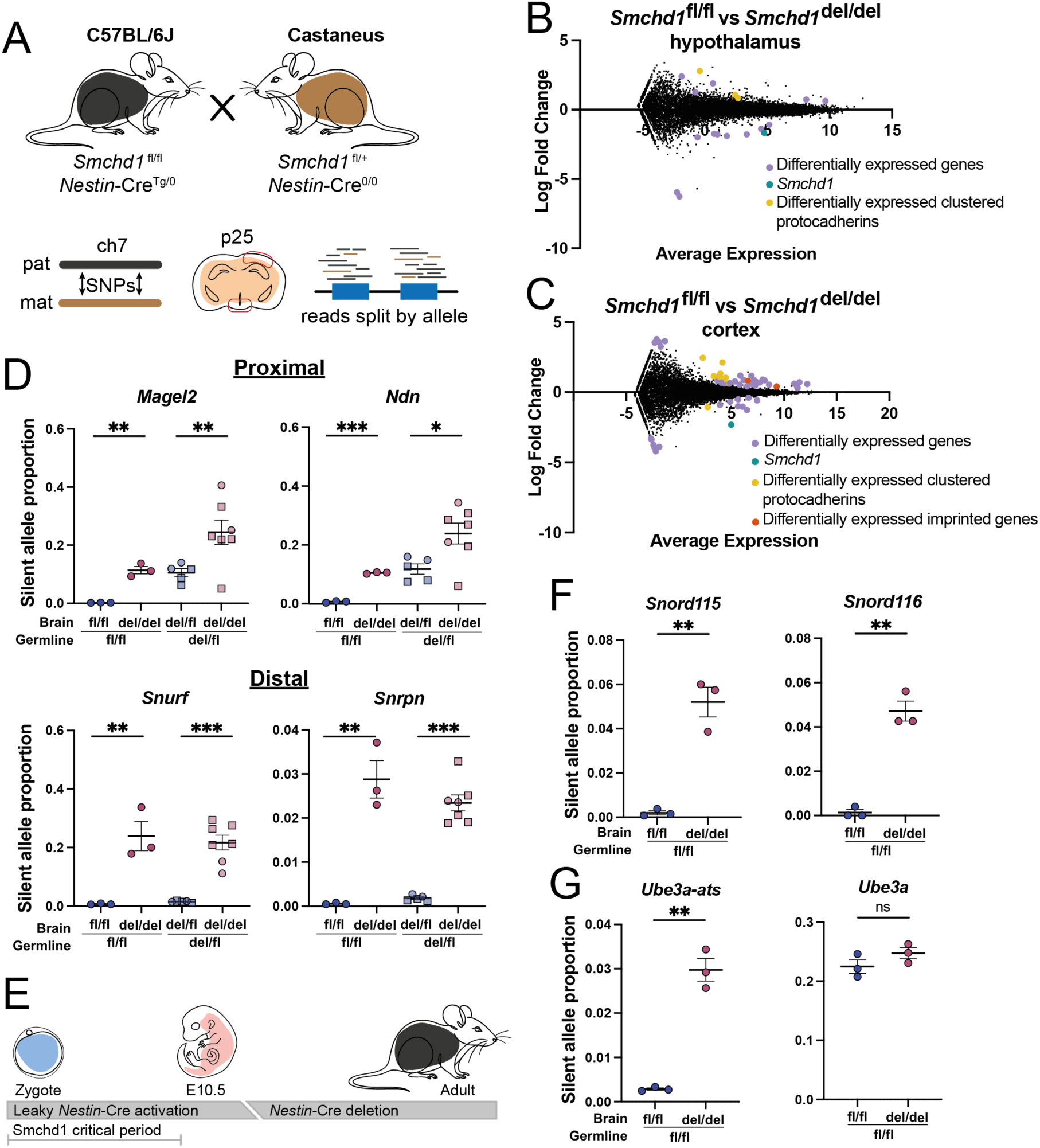
*Nestin*-Cre deletion of *Smchd1 in vivo* leads to activation of both proximal and distal maternal PWS cluster genes in the hypothalamus. **A)** Depiction of allele-specific sequencing methodology using strain-specific SNPs to determine parent-of-origin when mapping reads, red boxes indicate brain regions dissected for sequencing. **B-C)** MA plots showing differentially expressed genes between wildtype (*Smchd1*^fl/fl^, n=3) and *Smchd1* deleted (*Smchd1*^del/del^, n=3) female (**B**) hypothalamus (18 DE genes) and (**C**) cortex (56 DE genes) samples from RNA sequencing, FDR <0.05. **D)** Graphs depicting proportion of PWS gene expression from the imprinted maternal allele as a proportion of expression from the active paternal allele by RNA sequencing in hypothalamus of n=3 female mice. Circle data points are female, square are male, t-test with Bonferroni correction for multiple testing, ns = not significant * = FDR <0.05, ** = FDR <0.01, *** = FDR <0.001. **E)** Schematic depicting timing of *Nestin*-Cre deletion against the *Smchd1* critical developmental period, including overlap of stages with leaky Cre expression. **F-G)** Plots showing expression from the imprinted allele as a proportion of expression from the active allele by total RNA sequencing in the hypothalamus of n=3 female mice, t-test with Bonferroni correction for multiple testing, ns = not significant, ** = FDR <0.01.

Next, we performed allele-specific analysis by mapping the RNA sequencing reads to each parental genome, allowing us to consider the imprinted gene expression at the PWS locus. All PWS cluster genes were appropriately imprinted in the *Smchd1*^fl/fl^ hypothalamus samples; however, we observed significant increases in expression from the maternal allele at both proximal (*Magel2* and *Ndn*) and distal (*Snrpn* and *Snurf*) cluster genes when *Smchd1* was deleted (**Fig. 3D**). This was also seen to a lesser extent in the cortex, although not all PWS cluster genes were imprinted outside the hypothalamus (**Fig. S3**). Loss-of-imprinting at the distal end of the PWS cluster has not been previously reported in mice; however, nor has it been assessed using a sensitive allele-specific technique in a mature neural population. These data lend support to the hypothesis that SMCHD1 has brain-specific functionality at the PWS cluster.

Although *Nestin-*Cre provides efficient deletion in the brain, it is known to be leaky and can activate earlier than desired^68,66^ resulting in *Smchd1* deletion in the germline (**Fig. 3E)**. We defined leaky *Nestin*-Cre expression based on the *Smchd1* genotype in tail samples, where the presence of a *Smchd1*^del^ allele indicated deletion outside of the neural system, likely in the germline. In brain samples where leaky *Nestin*-Cre expression was present we observed similar levels of maternal *Magel2* and *Ndn* activation in *Smchd1^del^*^/fl^ hypothalamus to what was seen in the *Smchd1^del^*^/del^ samples (approximately 10% of active allele expression levels), *i.e.* heterozygous loss of *Smchd1* from the germline was equivalent in gene-activation to complete deletion in the brain with a wild-type level elsewhere in the animal. This was not seen for distal cluster genes in these samples, rather *Snurf* and *Snrpn* remained imprinted in the brain samples from germline *Smchd1*^del/fl^ animals (**Fig. 3D**). Furthermore, in the hypothalamus of animals that had germline *Smchd1* deletion followed by complete deletion in the brain, maternal *Magel2* and *Ndn* expression was elevated in comparison to the *Smchd1^del^*^/del^ animals without germline deletion (approximately 25% of active allele expression levels). Again, this was not seen for the distal PWS cluster genes, where an effect of *Smchd1* deletion required complete loss of *Smchd1* in the brain independent of a germline heterozygous deletion of *Smchd1* (**Fig. 3D**). These data suggest *Smchd1* deletion earlier in development results in more effective activation at the proximal end of the PWS cluster, but that control over distal imprinting in early development is maintained by mechanisms that do not depend on SMCHD1’s presence, consistent with our prior studies that focussed on such timepoints and also did not identify a role for SMCHD1 at the distal PWS cluster genes^51,58–62^.

Following the observation that *Smchd1* deletion results in maternal *Snurf* and *Snrpn* gene expression in the brain we performed stranded total RNA-sequencing using samples from *Smchd1*^del/del^ and *Smchd1*^fl/fl^ hypothalamus tissue with strain-specific SNPs. This allowed us to investigate expression of the *Snord115* and *Snord116* clusters, as well as *Ube3a* and *Ube3a-ats* as we could get both strand-specific and allele-specific information from this dataset. As expected, both *Snord* gene clusters exhibited increased expression levels from the maternal allele indicative of loss-of-imprinting at these loci (**Fig. 3F**). Our stranded analysis confirmed that *Smchd1* deletion also leads to an increase in maternal *Ube3a-ats* expression, however this had no impact on maternal *Ube3a* expression and is therefore unlikely to be detrimental to UBE3A expression levels in neurons (**Fig. 3G**), although the mechanism for this separation of expression warrants further enquiry.

### SMCHD1 has brain-specific binding patterns at the PWS cluster in mice

Given our new finding that in neural tissue SMCHD1 is involved in repressing the distal end of the PWS cluster as well as the proximal, we asked whether there were additional binding sites for SMCHD1 in mouse cortex samples. We performed ChIP-seq for GFP in *Smchd1*^GFP/GFP^ cortices; the *Smchd1*^GFP^ allele creates a SMCHD1-GFP fusion protein that we have shown retains normal function and allows us to take advantage of the GFP tag for immunoprecipitation^57,70^. We found 108,862 SMCHD1 binding sites genome-wide, including at known SMCHD1 targets and at new binding regions identified in this experiment (**Supplementary Table 5**). As expected, there was an abundance of peaks at known SMCHD1 targets such as 666 sites across the clustered protocadherins (**Fig. 4A, Supplementary Table 6**) and 61,814 on the X chromosome (**Fig. 4B, Supplementary Table 7**), likely representing the inactive X as these were all female samples. To query whether binding was required for silencing, we utilised our RNA-seq dataset to investigate SMCHD1 binding at regions containing differentially expressed genes between *Smchd1* wild-type and *Smchd1* deleted cortices, to match the tissue used for ChIP. We noticed a few examples of differentially expressed genes falling within highly SMCHD1-enriched gene-clusters. *ZfpS45* was differentially expressed in our cortical RNA-seq dataset and is located within a cluster of zinc-finger proteins that has 1,412 SMCHD1 binding sites (**Fig. 4C, Supplementary Table 8**). Notably two of the differentially expressed genes, *Hbb-b2* and *Syt9* were each within one of two large adjacent clusters of olfactory receptor genes spanning a combined 6.4 Mbp, across which there are 297 SMCHD1 peaks (**Fig. S4A, Supplementary Table 9**). Additionally, there was one differentially expressed gene (*E2f8*) within 150kb of another large SMCHD1 bound gene cluster, the Mrgpra cluster of sensory neuron receptor genes at which there are 734 SMCHD1 peaks (**Fig. S4B, Supplementary Table 10**). Overall, of the 56 genes that are differentially expressed 31 could be considered as SMCHD1 targets with SMCHD1 peaks located within 150 kb of the gene; of these only 23 contain peaks within the gene and proximal promoter (**Supplementary Table 11**). However, there are only ever one or two differentially expressed genes within megabase-long highly SMCHD1 enriched regions, which indicates that expression of the vast majority of SMCHD1 targets remain unaffected when SMCHD1 is deleted from mid-gestation in the brain.

**Figure 4:**
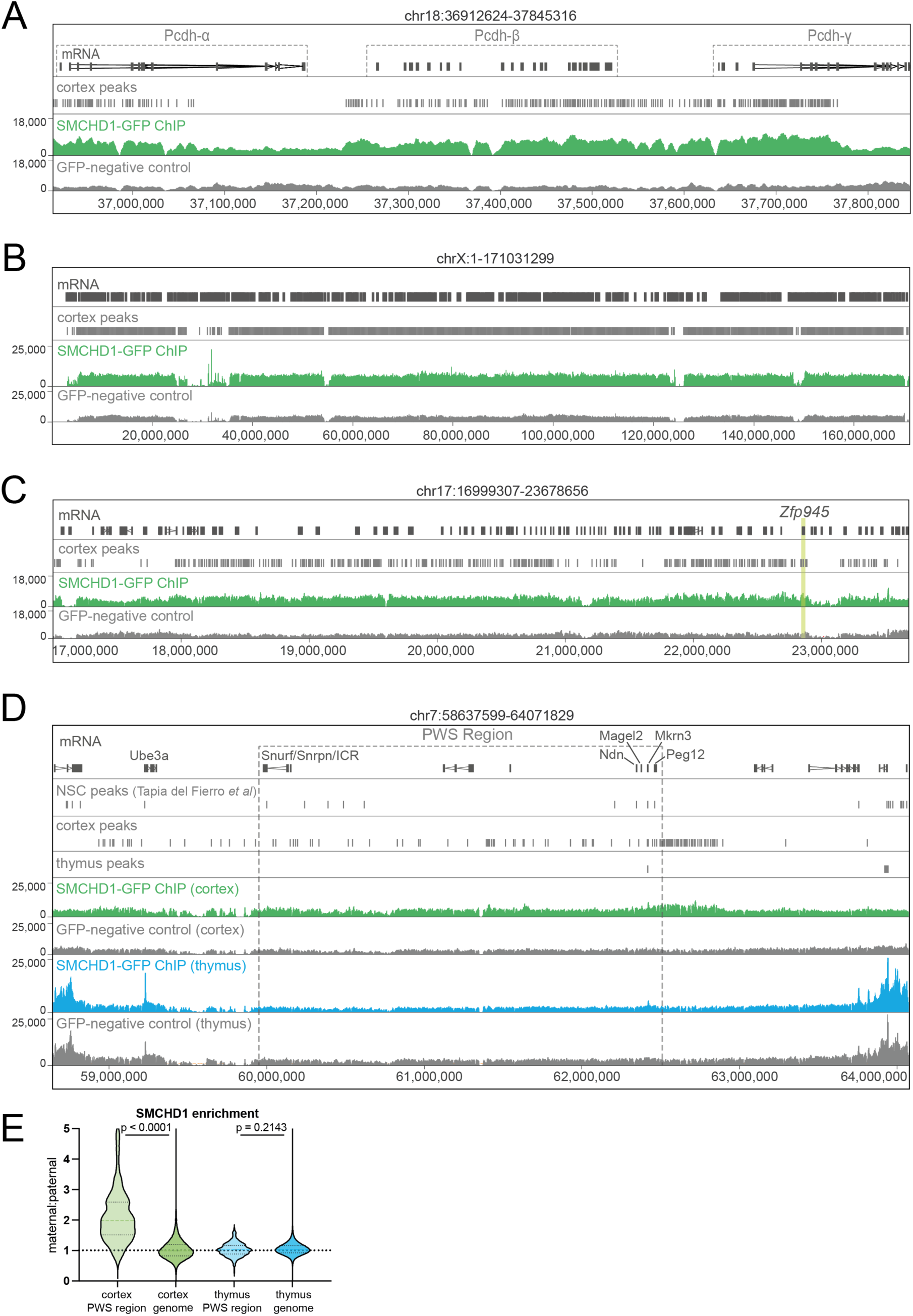
SMCHD1 binding is enriched at the PWS cluster in the brain. Genome browser ChIP-seq tracks for SMCHD1-GFP ChIP-seq showing peaks at **A)** the clustered protocadherins locus, **B)** the X chromosome, **C)** the zinc-finger protein cluster with *ZfpS45* highlighted as differentially expressed between *Smchd1*^del/del^ and *Smchd1*^fl/fl^ cortex samples by RNA-seq. n=2 for each sample. **D)** Genome browser ChIP-seq tracks for SMCHD1-GFP ChIP-seq at the PWS cluster and flanking non-imprinted regions showing peaks from previously published in-vitro data (NSC peaks^70^), and peaks for cortex tissue samples or thymus tissue samples (n=2 for each). **E)** Graph plotting the ratio of SMCHD1 enrichment on the maternal versus paternal alleles from SMCHD1-GFP ChIP-seq using 10 kb bins to section the genome, data from the PWS region and sex chromosomes were excluded from the genome wide data. Coloured dashed lines at mean, n=2 for each tissue. Welch’s t-test significance = p <0.05.

At the PWS locus we discovered additional SMCHD1 binding sites in the cortex samples compared both with previously published SMCHD1 ChIP-seq datasets^51,58,70^ and with matched thymus samples as a non-neural example tissue (**Fig. 4D, Supplementary Tables 12-13**). As prior neural datasets have come from embryonic cortical neural stem cells^51,58,70^ we hypothesise this increase in SMCHD1 binding at the PWS cluster to be specific to mature neurons where expression of PWS imprinted genes is most prominent. We further hypothesised that neural SMCHD1 binding at the PWS cluster would be predominantly on the silent maternal allele, given that SMCHD1 removal leads to activation of maternal PWS genes. To test these hypotheses, we performed allele-specific ChIP-seq using the same *Smchd1*^GFP/GFP^ animals (C57BL/6J background) crossed with a Castaneus strain to again exploit SNPs between alleles. Contrary to expectations we observed reads on both maternal and paternal alleles at the PWS cluster. Nevertheless, using a sliding-window based approach we saw an increase in the ratio of reads coming from the maternal allele versus the paternal allele at the PWS locus compared with elsewhere in the genome in cortex samples; this was not seen in the thymus samples where parent-of-origin ratios remained around 1:1 (**Fig. 4E**). Taken together these data suggest that while SMCHD1 is not required for the maintenance of silencing at the majority of its binding sites across the genome it is required at the PWS cluster, thereby affirming SMCHD1’s importance, particularly on the maternal allele at the region.

### Smchd1 deletion in a mouse model of PWS can ameliorate some disease phenotypes in vivo

We sought to understand whether the maternal PWS gene-activation achieved with neural *Smchd1* deletion *in vivo* is sufficient to improve PWS phenotypes in a *Magel2*-null mouse model of PWS. This mouse model utilises the *Magel2*-*LacZ* knockin-knockout strain^66^ used as a reporter in earlier experiments (**Fig. 2**); however, the disease-model mice are rendered *Magel2*-null by having the *LacZ* knockin on the paternal i.e. active allele. This results in mice with no active paternal copy of *Magel2* and a remaining silenced maternal copy (**Fig. 2B**) akin to PWS patients. Aligned with symptoms of PWS in human patients the *Magel2*-null mice in this model have been reported to have various muscular, motor and behavioural deficiencies^66,71–74^.

Due to the high variance observed in behavioural studies, and therefore the requirement for high sample numbers^75^, we created two separate experimental cohorts. All data was analysed collectively for maximum statistical power with sex and cohort as variables. Here we only report data with no significant cohort effect and combine sexes only where there was no significant sex-specific effect. One important caveat regarding these cohorts arises from the aforementioned leaky *Nestin*-Cre. Across our two large cohorts, animals that were *Smchd1*^fl/fl^ and *Nestin*-Cre transgenic were not found. Rather, all *Nestin*-Cre transgenic offspring were either *Smchd1*^fl/+^ or *Smchd1*^fl/del^ by ear-clip genotyping, i.e. a non-neural tissue where *Nestin*-Cre should not delete, suggesting *Smchd1* deletion by leaky Cre in the male germline. Therefore, by necessity all control mice within these cohorts were already *Smchd1*^del/+^ in the brain rather than wild-type for *Smchd1*.

Through our panel of tests we observed PWS phenotypes in our cohorts that match published data for this model. In *Magel2*-null mice (*Smchd1*^del/+^*Magel2*^patKI/+^*Nestin*-Cre^Tg/0^), henceforth called PWS control mice, both males and females had increased body weight with age, decreased grip strength, and impaired motor coordination compared to *Magel2* replete (*Smchd1*^del/+^*Magel2*^+/+^*Nestin*-Cre^Tg/0^), henceforth referred to as wild-type controls (**Fig. 5A-C, S5A-C)**. Sex-specific effects of *Smchd1* deletion were apparent in our data. No improvements were seen in male *Magel2*-null mice with a *Smchd1* deletion in the brain (*Smchd1*^del/del^*Magel2*^patKI/+^*Nestin*-Cre^Tg/0^), henceforth referred to as our test animals, for body weight (**Fig. S5A**) nor grip strength (**Fig. S5B**); however, they did perform slightly better at lower speeds in the motor coordination test (**Fig. S5C**). For all three of these phenotypes the female test animals were not significantly different from wild-type indicating phenotypic improvement in PWS mice upon *Smchd1* deletion (**Fig. 5A-C**). Further, we noticed the female weight data for the test group was bimodal, with 45% of the test animals clustering within the wild-type control group and the remaining 55% clustering within the PWS control group (**Fig. 5A**). This suggests that instead of an overall marginal improvement in the weight phenotype we observe a rescue in 45% of test animals. We asked whether the animals with a phenotypic weight rescue performed less like PWS controls in our other tests. For grip strength, noting that it is weight-normalised data, we observed a statistically significant rescue in this subset when tested against PWS controls at 22 weeks of age (**Fig. 5B**). In the muscle coordination data, which is more variable, there was no statistically significant rescue; however, three of the five test animals from our weight-rescued subset performed the best across all speeds (**Fig. S5D**). Together these data indicate that by deleting *Smchd1* in the brain we can rescue muscular and body composition PWS phenotypes in female mice in this model.

**Figure 5:**
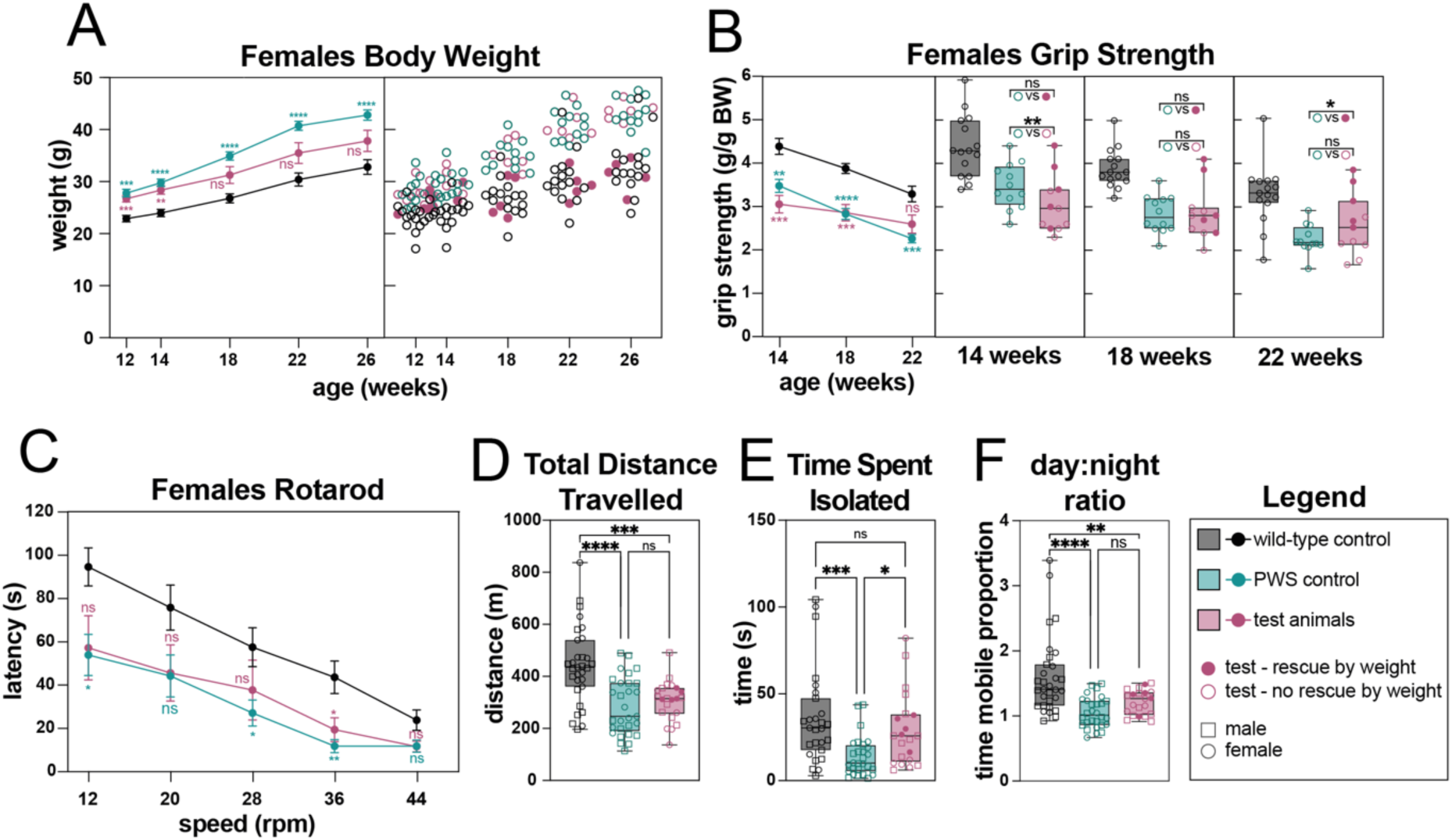
*Nestin*-Cre deletion of *Smchd1* can ameliorate disease phenotypes in a *Magel2*-null mouse model of PWS. n=11-16 per genotype across two cohorts, teal asterisks compare PWS control and wild-type control groups, pink asterisks compare wild-type control and test animal groups. **A)** Total body weight in female mice, left panel shows averaged data, right panel shows mice as individual points displaying bimodal distribution of test animals. **B)** Grip strength measurements normalised to body weight in female mice, left panel has averaged data for all time points tested, right panels plot individuals at each time point. **C)** Graph plotting average latency to fall from two rounds of testing for motor coordination on a rotarod in female mice. **D-F)** Plots from a home-cage observation period of >44h including both male and female mice across two cohorts and showing (**D**) Total distance travelled. (**E**) Average time spent isolated from all other cage-mates per hour from two combined 10h active periods, each cohort normalised to average of wildtype for statistical analysis. (**F**) The ratio of time spent mobile in the active versus inactive periods. Genotypes in the brain are wild-type control = *Smchd1*^del/^*^ff^Magel2*^+/+^, PWS control = *Smchd1*^del/^*^ff^Magel2*^patKI/+^, test animals = *Smchd1*^del/del^*Magel2*^patKI/+^, error bars represent ±SEM, two-way ANOVA Tukey’s multiple comparison **(A-C)** or one-way ANOVA Šidák’s multiple comparison **(D-F)** adjusted p-values ns = not significant, * = p <0.05, ** = p <0.01, *** = p <0.001, **** = p <0.0001.

To assess cognitive and social phenotypes without introducing undue stress responses in our mice we observed the cohorts using the Actual Analytics Home Cage Analyser^76^. We recorded observations from two active periods (night) and one inactive period (day) and combined data from all mice across both cohorts. In line with the muscular defects described above between PWS control and wild-type control groups we detected a significant decrease in the total distance travelled during this time by the PWS control mice. This was true for both sexes; however, unlike other muscular/motor phenotypes this was not improved with *Smchd1* deletion in either males or females (**Fig. 5D**). Using data from both active periods we found PWS control mice engage in significantly less time isolated from cage mates compared with wild-type mice. This phenotype was rescued by *Smchd1* deletion in both males and females as evidenced by a statistically significant difference between the PWS control group and our group of test animals, with no statistical difference found between test and wild-type groups (**Fig. 5E**).

PWS patients commonly suffer from sleep disorders^8,77,78^. To investigate circadian rhythm defects in our cohorts we compared activity ratios during active versus inactive periods from the home cage observation data. We found that there was a significant decrease in the day:night activity ratio when looking at the average amount of time the PWS control mice spent mobile compared with the wild-type controls, again for both sexes. This difference is reduced but remains between the wild-type and test groups (**Fig. 5F**), which could indicate improvement in this phenotype resulting from *Smchd1* deletion. These data suggest increased power from considerably more replicates would be required to interrogate this further.

Within this cohort we also obtained data from *Magel2* replete but *Smchd1*-null animals (*Smchd1*^del/del^*Magel2*^+/+^*Nestin*-Cre^Tg/0^) and were able to compare these to our wild-type mice to assess the effect of *Smchd1* deletion independent of any PWS phenotype effects. We observed a significant increase in body weight for male mice throughout life, and a transient decrease in grip strength for male mice that was only apparent at 18 weeks of age **(Fig. S5E-F)**. For all other measures in male mice, and for all measures in female mice we saw no significant effects of *Smchd1* deletion on mice with functional *Magel2* **(Fig. S5E-J)**.

Taken together our functional studies demonstrate that *Smchd1* deletion can be beneficial by improving disease-relevant phenotypes in a mouse model of PWS. It further highlights that this approach is likely to be safe since we observed no detrimental effects in our test animals, and the *Smchd1* deleted animals had normal survival.

## DISCUSSION

This body of work aimed to examine the potential of SMCHD1 as a target for gene-activation therapeutics in the treatment of PWS in light of its prior known role in silencing maternal PWS cluster genes in mice. Importantly, here we discovered SMCHD1 to likewise contribute to maternal PWS gene silencing in human cells, phenocopying findings in mice. Using PWS patient-derived iPSC models we found that not only does removal of SMCHD1 lead to maternal loss of imprinting in early development, but that this can also be seen when SMCHD1 is removed in a more committed lineage, more akin to timing of therapeutic interventions. These data provide evidence for the potential clinical utility of such an approach for treating the genetic cause of PWS in patients.

We next addressed the questions of efficacy and safety using mouse models to assess both functional and off-target effects of targeting SMCHD1 *in vivo*. Following confirmation that *Smchd1* deletion *in vivo* does indeed cause activation of maternal PWS cluster genes in the brain we looked more closely into the importance of timing for this intervention. Proximal PWS cluster genes *Magel2* and *Ndn* are more SMCHD1 sensitive in that the level of gene-activation observed is dependent on timing of *Smchd1* deletion. When deleted earlier in development there was a more marked increase in maternal *Magel2* and *Ndn*, whereas for distal PWS cluster genes *Snurf* and *Snrpn* the amount of maternal gene-activation remained constant irrespective of when *Smchd1* was removed. The sensitivity of proximal PWS genes was also apparent in the case of heterozygous loss of *Smchd1*, whereas distal genes were not affected in this scenario; this might suggest that later intervention could be more selective.

Furthermore, we discovered an enhanced role for SMCHD1 at the PWS cluster in neurons, which proved true in both mouse and human. Previous data from non-neural or immature neural populations showed a role for SMCHD1 at only the proximal end of the PWS cluster^51,58–62^. We revealed SMCHD1’s role at the distal end of the locus by looking in PWS patient-derived iPSCs differentiated down a neural pathway, as well as in mouse brain tissue containing mature neurons. Relatedly, we discovered extra enrichment of SMCHD1 binding in brain tissue samples compared with both a non-neural control tissue and our previous findings in neural progenitor cells^51,58,70^. This serves to support the concept of neural-enhanced functionality for SMCHD1 at the PWS locus. SMCHD1 binding at the PWS locus also seems to be enriched on the imprinted maternal allele, which is not the case elsewhere in the genome. Genome-wide we found few differentially expressed genes by RNA-seq as a result of *Smchd1* deletion; most of these were the singular differentially expressed exception within highly SMCHD1 enriched regions by ChIP-seq, suggesting that loss of SMCHD1 mid-gestation does not lead to loss of silencing for the majority of target genes. In conjunction with timing of deletion being a selective factor, these data further support the potential safety of SMCHD1 targeting as its effect appears quite specific to mature neurons at the PWS cluster with low impact elsewhere around the genome.

We wanted to address the question of safety in terms of a potential effect on maternal *Ube3a* gene expression specifically, particularly in light of the discovery that *Smchd1* deletion can increase maternal snoRNA expression in neural populations at similar levels to what was seen for *Snrpn*. We found significant upregulation of maternal *Ube3a-ats*, potentially as a consequence of activating transcription at the *Snurf-Snrpn* shared promoter. Interestingly, we did not see any change in maternal *Ube3a* gene expression suggesting that the degree of imprinting loss at *Ube3a-ats* was not sufficient to disrupt *Ube3a* expression as is the case on the paternal allele. This suggests no risk of inadvertent Angelman Syndrome while treating PWS; however, if methods to boost activation beyond what we can currently achieve were to be employed this would need to be reassessed.

Our functional studies in a mouse model of PWS were able to recapitulate some muscular and behavioural PWS phenotypes *in vivo*. Due to the highly variable nature of analysing PWS-related phenotypes on a whole-organism scale^75^ we took a conservative approach and only included data significant across two separate cohorts of animals. There were also some outstanding sex-specific differences, which is a common feature of these types of studies and can further reduce statistical power. Nevertheless, we were able to detect improvements in weight, grip strength and motor coordination phenotypes in female PWS mice that had *Smchd1* deleted. Remarkably, we saw full rescue of a social-behavioural phenotype, time spent in isolation, when able to combine both cohorts and sexes. As noted, the PWS mice in our cohorts were already heterozygous for *Smchd1* due to practical limitations of the study. Therefore, it is likely we began with diminished capacity to detect PWS phenotypes as we could potentially be masking the severity of these with already reduced *Smchd1* presence. If our ability to detect phenotypes is lessened, there would likewise be reduced power to detect improvements and/or rescue. Considering the safety perspective, if SMCHD1 targeting were to ever be applied in a therapeutic setting, the *Smchd1* deleted mice in these cohorts remained healthy into adulthood with a small increase in male weights and a transient decrease in male grip strength being the only observed effects of *Smchd1* deletion. As a whole, we believe these data to be an encouraging step towards confirming the clinical utility of SMCHD1 as a target for epigenetic therapy to treat PWS.

Future work using non-genetic approaches that directly target SMCHD1 will be beneficial as it will eliminate problems associated with *Nestin*-Cre, enable more precisely timed treatments and more closely mimic clinical applications. The novel discovery of SMCHD1’s role across both the proximal and distal ends of the PWS locus means other mouse models would also be useful for future investigations. For instance, whole-locus disruption models such as the imprinting centre deletion^76^, while neonatal lethal, could be useful to simply test for increased survivability with SMCHD1 removal.

As mentioned, most research into gene-activation approaches for PWS has been focused on the distal end of the locus. We have found proximal PWS cluster genes to be particularly SMCHD1 sensitive, as well as a neural-enhanced role for SMCHD1 encompassing the entire locus. The gene-activation that we attain by targeting SMCHD1 alone does not reach paternal PWS gene expression levels, confirming that other factors are also relevant to imprinting maintenance at the cluster^15,24–28^. More in-depth investigation into SMCHD1’s mechanism of maternal gene silencing at the human PWS cluster will aid in determining which pathways may synergise and contribute to increased understanding of the overwhelmingly complex epigenetic control in the region. Thus, our research here is a vital stepping stone, which could lead to advancement of prospects for epigenetic therapy to treat PWS.

## MATERIALS AND METHODS

### Cell culture

HEK293T cells were cultured in DMEM (Gibco) supplemented with 10% FBS (Life Technologies) at 37°C in a humidified cell culture incubator with 10% (v/v) CO2. Cells were passaged before reaching 85% confluency by dissociation with 0.5% Trypsin-EDTA (Life Technologies).

iPSCs were purchased from University of Connecticut Stem Cell Core (see **Supplementary Table 2** for details) and cultured in mTesR™1 medium supplemented 1x antibiotic-antimycotic solution (Sigma); on plates coated with 0.12-0.18mg/ml Geltrex (Gibco) in Knockout DMEM (Gibco) at 37°C in a humidified cell culture incubator with 10% (v/v) CO2. Before reaching 80% confluency cells were passaged as aggregates by dissociation with Versene Solution (Gibco) and gentle lifting with a cell scraper. Aggregates were broken up using gentle trituration with a wide-bore pipette tip and then were transferred to a new Geltrex-coated well. Media was supplemented with 2µM Thiazovivin (Axon) overnight immediately after passaging.

NPCs were cultured in neural media (described below) supplemented with 10ng/ml FGF (Stem Cell Technologies) and 10ng/ml EGF (Stem Cell Technologies) at 37°C in a humidified cell culture incubator with 10% (v/v) CO2.

### Lentivirus-mediated CRISPR-Cas9 knockout of SMCHD1 in HEK293Ts and PWS patient-derived iPSCs

gRNAs were cloned into lentiGuide-Puro (addgene plasmid #52963) and pFgH1UTPuro (addgene plasmid #70183 with the GFP reporter replaced by puromycin resistance).

HEK293T cells were cultured as described above. HEK293T cells at 80% confluency were transfected via calcium phosphate mediated transfection with the psPAX packaging vector, the VSVg envelope vector and one of the desired plasmids containing either SMCHD1 guides or Cas9 (addgene plasmid #52962) in the ratio 2.5:1:1.5. Transfection cocktails made up of plasmid DNA and 250mM CaCl2 were precipitated in HBS solution and then added in a dropwise fashion to the plated HEK293T cells and left overnight at 37°C. Media was replaced after 16h and supernatant was harvested 24 hours later through a 0.45µM filter and concentrated with Poly(ethylene glycol) (PEG). Filtered supernatant containing lentivirus was transferred to a tube containing PEG-8000 solution [PEG-8000 40% (w/v), 1.2M NaCl in 1x phosphate-buffered saline (PBS), pH 7.0-7.2] in a 3:1 volume ratio. Solution was mixed by shaking for 60 seconds and then incubated at 4°C for >16h with constant rocking. Supernatant was removed following centrifugation at 1600xg for 60 minutes and viral pellet was thoroughly resuspended in 1/10 the original volume of serum-free culture media by gentle trituration. Concentrated virus was aliquoted and stored at -80°C until use.

1:50 - 1:100 dilution of concentrated virus was added to the media of HEK293T cells or iPSCs at approximately 50% confluency along with 4µg/ml polybrene (polybrene excluded for iPSCs) (Sigma-Aldrich). Media was changed after 24 hours to media containing the appropriate antibiotic for selection. Blasticidin (Thermo Fisher) used at 7µg/ml (HEK293Ts) or 5µg/ml (iPSCs) for >5 days, puromycin (Sigma-Aldrich) used at 2µg/ml (HEK293Ts) or 0.5µg/ml (iPSCs) for > 24h. 1x CloneR™2 (Stem Cell Technologies) was added to iPSC transductions and to seeded clonal cell lines. Clonal cell lines were established by limited dilution and editing efficiency was assessed by MiSeq DNA sequencing (Illumina) following PCR amplification of the targeted region and secondary PCR with overhang sequences (see **Supplementary Table 14** for primer sequences).

### Immunofluorescence microscopy

Immunofluorescence was performed the same for all cell types. Cells were grown on glass coverslips or in chamber slides and washed with PBS prior to starting. Cells were fixed in 4% (w/v) paraformaldehyde/PBS for 10 minutes at room temperature followed by 3x PBS washes. Permeabilisation was done for 5 minutes on ice in cold 0.5% (v/v) Triton X-100/PBS followed by a further 3x PBS washes. Cells were blocked in 1% (w/v) bovine serum albumen (BSA) (Gibco) for 30 minutes at room temperature. Primary antibodies were diluted 1:100 (SMCHD1 in house monoclonal clone 2B8) or 1:50 (SNRPN PA5-72858 ThermoFisher) in 1% (w/v) BSA and incubated for 1 hour at room temperature in a humidified chamber followed by 3x PBS washes. Secondary antibodies (AlexaFluor 488 donkey anti-rabbit IgG A21206; AlexaFluor 555 goat anti-rat IgG A21434) were diluted 1:500 in 1% (w/v) BSA and incubated for a further hour at room temperature. Cells were washed 3x with PBS, once for 1 minute with 2mg/ml DAPI nuclear stain (Thermo Fisher) diluted 1:10,000 in PBS, 3x further PBS washes, coverslips mounted on glass slides using VECTASHIELD®Vibrance™ antifade mounting medium (Vector Laboratories). Cells were visualised using an LSM 880 Upright (Zeiss) confocal microscope, analysis was done using the open-source ImageJ distribution package FIJI and/or manual scoring. Prism software (GraphPad) was used for statistical analysis and to plot data.

### Differentiation of iPSCs and NPC culture

iPSCs differentiated either using STEMdiff™ SMADi Neural Induction Kit (Stem Cell Technologies as per the manufacturer’s protocol), or using dual SMAD inhibition as follows: iPSCs singularised by incubating with StemPro™Accutase™ (Gibco). Cells seeded in mTesR™1 medium containing 2µM Thiazovivin (Axon) at 800K cells/cm^2^ onto a Geltrex (Gibco) coated plate and left to settle overnight. Cultured for 9 days in neural media [1:1 DMEM/F12 + Glutamax (Gibco):Neurobasal medium - Glutamax (Gibco), 0.5x B27 supplement (Gibco), 0.5x N2 supplement (Gibco), 0.5x Glutamax (Gibco), 0.5x NEAA (Gibco), 50µM β-mercaptoethanol, 1x ITS-A (Gibco), 1x antibiotic-antimycotic solution (Sigma)]; supplemented with fresh 10µM SB431542 (Stem Cell Technologies) and 100nm LDN 193189 (Selleckchem) daily. On day 10 cells were passaged with versene solution (Gibco) 1:2 into a Geltrex (Gibco) coated plate in neural media containing 10µM SB431542 (Stem Cell Technologies), 100nm LDN 193189 (Selleckchem), and 2µM Thiazovivin (Axon) overnight. Media changed to neural media only for 1 day, and then to neural media supplemented with 10ng/ml FGF (Stem Cell Technologies) and 10ng/ml EGF (Stem Cell Technologies) for 4 days. Cells passaged 1:6 with versene solution (Gibco) onto Geltrex (Gibco) and cultured in neural media for 7 days. Neural progenitor cells were singularised with StemPro™Accutase™ (Gibco), passaged 1:6 onto Geltrex (Gibco) and maintained in neural media supplemented with 10ng/ml FGF (Stem Cell Technologies) and 10ng/ml EGF (Stem Cell Technologies).

### qRT-PCR

RNA was extracted using the Quick-RNA Miniprep Kit (Zymo) as per manufacturer’s instructions altered only to increase incubation time with DNaseI to 45-60 minutes. cDNA was synthesised from 500-1000ng of RNA in 11µl nuclease free water. 2µM (final) Oligo(dT) primer (Promega) and 0.4µM (final) dNTP mix (Promega) was added to the RNA and incubated at 65°C for 5 minutes. Added to this was 1x RT buffer (Thermo Fisher), 4mM MgCl2 (Promega), 8mM DTT (Thermo Fisher), 40 units RNaseIN (Thermo Fisher), 200 units Superscript III Reverse Transcriptase (Thermo Fisher). Control reactions for each sample containing no reverse transcriptase were also created at this point and included in every qRT-PCR experiment. Reactions were then incubated at 50°C for 50 minutes followed by 5 minutes at 85°C. cDNA was used at either 1:5 (HEK293Ts) or 1:3 (all other cells) dilution for the RT-qPCR reaction comprising 1/5 diluted cDNA template, 0.5µM forward and reverse primers (see **Supplementary Table 14**), 1x FastStart Universal SYBR Green Master Mix (Roche) or 1x SensiMix™ SYBR® Hi-ROX (Bioline), and nuclease free water. PCR reactions were performed using either the LightCycler® 480 Real-Time PCR Instrument (Roche) or the QuantStudio™ 12K Flex Real-Time PCR System (Thermo Fisher) with cycling conditions of 50° C for 2 minutes, 95°C for 10 minutes; 40x cycles of, 95°C for 15 seconds, 60°C for 1 minute. Cycle thresholds (Ct) were calculated using the instrument software and relative mRNA expression levels were calculated by double delta Ct analysis using GUS as a housekeeping gene. Prism software (GraphPad) was used for statistical analysis and to plot data.

### *Smchd1* deletion by nucleofection of CRISPR ribonucleoproteins

50µM gRNA per nucleofection reaction was prepared by incubating Alt-R® CRISPR-Cas9 tracrRNA (IDT) and SMCHD1 crRNA (see **Supplementary Table 14**) (100µM in sterile Tris-EDTA buffer) in a 1:1 ratio, heating to 95°C for 5 minutes and cooling to room temperature. 10µg TrueCut Cas9 protein (ThermoFisher) was added and incubated for 20 minutes at room temperature. 1 million NPCs per nucleofection reaction were singularised using StemPro™Accutase™ (Gibco), quenched in neural media and pelleted at 100xg for 10 minutes. Cells were washed in PBS and pelleted as previous. Cells resuspended in P3 Primary Cell Nucleofector® Solution plus Supplement 1 from the Amaxa P3 primary cell nucleofector kit (Lonza Bioscience) and combined with the prepared RNP complex and 4.8µM Alt-R® Cas9 Electroporation Enhancer (IDT). Each reaction was pipetted into one well of a Nucleocuvette® (Lonza Bioscience) and nucleofected using pulse code CA-137 on the Lonza 4D-Nucleofector®. Electroporated samples were transferred to prewarmed neural media containing 2µM Thiazovivin (Axon). Media changed after 24 hours and then cells were cultured using standard methods for 7 days before harvest and assay.

### Depletion of SMCHD1 in NPCs via exon-skipping using ASOs

On day 0 a confluent well of NPCs was treated with 1µM ASO, either targeting SMCHD1, or targeting Dystrophin, which is not expressed in NPCs and was thus used as a non-targeting control (see **Supplementary Table 14** for ASO sequences). ASO was added directly to the culture media, mixed gently by trituration and incubated at 37°C for 2 hours after which the media was replaced with fresh neural media. This was repeated until day 7 with the inclusion of a single PBS wash following treatment on days 5 and 6 to remove dead cells. Cells were harvested on day 7 and RNA was purified using the Quick-RNA miniprep kit (Zymo) as per manufacturer’s instructions. Exon skipping was assessed using cDNA synthesised as described above from 300ng RNA in nuclease free water. cDNA was used 1:20 as the template in the RT-PCR reactions comprising 1x GoTaq® Green Master Mix (Promega), 0.5µM of each forward and reverse primer (see **Supplementary Table 14**) in nuclease-free water. Cycling conditions were 94°C for 5 minutes; 35x cycles of, 94°C for 30 seconds, 55°C for 30 seconds, 72°C for 1 minute; 72°C for 7 minutes. PCR products were resolved by gel electrophoresis on a 2% agarose gel.

### Mouse strains and genotyping

All mice were bred and maintained as per standard husbandry procedures approved by the WEHI animal ethics committee under animal ethics numbers 2020.050 then 2023.033, all behavioural procedures were performed under animal ethics number 2020.058.

Rosa26^CreERT2^ mice carry a knockin allele to express CreERT2 under the transcriptional control of the ubiquitously expressed *Rosa26* locus^67^. These mice are on a C57BL/6J strain background.

*Magel2*-*LacZ* knockout knockin mice have the entire *Magel2* reading frame replaced by a *LacZ*-neo cassette with the endogenous *Magel2* promoter driving *LacZ* expression^66^. These mice are on a C57BL/6J strain background. Mice were obtained from The Jackson Laboratory (Jax stock #009062)

*Nestin*-Cre transgenic mice express Cre recombinase controlled by the *Nestin* promoter and nervous-system-specific enhancer to express Cre in glial and neuronal cell precursors from embryonic day 10.5^63^. These mice are on a C57BL/6J strain background.

*Smchd1^fl^* mice contain a floxed allele of *Smchd1* with Lox-P sites flanking exon 5 to enable specific Cre recombinase mediated deletion of *Smchd1*^64^. These mice are on a C57BL/6J strain background, except for allele-specific studies, where we had backcrossed the mice for 5 generations onto the Castaneus background.

*Smchd1^GFP^* mice have a GFP knockin at the *Smchd1* locus immediately preceding the stop codon to create a Smchd1-GFP fusion protein^58^. These mice are on a C57BL/6J strain background.

Castaneus (Cast) mice founders were trapped in a grain warehouse in Thailand in the 1970s^80,81^. The mice used in this project came from The Jackson Laboratory stock #000928.

Samples for genotyping were lysed overnight at 55°C in Direct PCR Lysis reagent (Viagen) containing 200µg/ml Proteinase K (Roche), the Proteinase K was heat inactivated at 85°C for 45 minutes. The resulting lysate was used as the template to determine genotypes by PCR. Each PCR reaction contained 1:20 DNA template, 1x GoTaq? Green Master Mix (Promega), 0.5µM of each forward and reverse primer (see **Supplementary Table 14**) in nuclease-free water. Cycling conditions were 94°C for 5 minutes; 35x cycles of, 94°C for 30 seconds, 60°C for 30 seconds, 72°C for 1 minute; 72°C for 7 minutes, 12°C hold. PCR products were resolved by gel electrophoresis on a 1.8% agarose gel.

### Tamoxifen treatments in neonates

All tamoxifen treatments were approved by the WEHI animal ethics committee under animal ethics number 2020.049. Tamoxifen was administered by a mouse technician at either 50µg/g body weight, 5x daily doses (postnatal day 9-13) by oral gavage or 50µg 2x daily doses (postnatal day 1-2) by intragastric injection. All administrations were performed according to WEHI standard operating procedures. Mice were monitored twice on the day of treatment and daily thereafter. Where necessary and by WEHI veterinarian approval pups were supplemented with Di-Vetelact® nutritional milk. Mice were considered at endpoint after two consecutive days of weight loss.

### Cryosectioning and X-gal staining of mouse brains

Brains from *Magel2*-*LacZ* mice were dissected directly into Tissue-Tek® O.C.T Compound (Sakura), frozen on dry ice and stored at -80°C until cryosectioning. Sections were cryosectioned on the HM525 Crysotat (Thermo Scientific), chamber temperature -15°C, specimen temperature -18°C, 12µM slice thickness, and mounted onto Fisherbrand™Superfrost™Plus Microscope Slides (Fisher Scientific).

β-galactosidase activity was detected by X-gal (5-bromo-4-chloro-3-indolyl-beta-D-galacto-pyranoside) staining. Sections were fixed for 15 minutes at room temperature in fix buffer [0.1M phosphate buffer (3.74g monobasic sodium phosphate (Sigma-Aldrich), 10.35g dibasic sodium phosphate (Sigma-Aldrich), in 1L H2O (pH7.3)), 5mM EGTA (Sigma-Aldrich), 2mM MgCl_2_ (Sigma-Aldrich), 0.2% (v/v) glutaraldehyde (Sigma-Aldrich)]. Sections were washed twice in wash buffer for 5 minutes each wash [0.1M phosphate buffer, 2mM MgCl2 (Sigma-Aldrich), 0.01% (v/v) deoxycholate (Sigma-Aldrich), 0.02% (v/v) NP-40 (Sigma-Aldrich)]. Sections were incubated at 37°C for 1.5-3 hours in X-gal staining buffer [0.1M phosphate buffer, 2mM MgCl_2_ (Sigma-Aldrich), 5mM potassium ferrocyanide (Sigma-Aldrich), 5mM potassium ferricyanide (Sigma-Aldrich), 0.01% (v/v) deoxycholate (Sigma-Aldrich), 0.02% (v/v) NP-40 (Sigma-Aldrich), 1mg/ml X-gal (Sigma-Aldrich)]. Stained slides were counterstained with Nuclear Fast Red and imaged on the Vectra Polaris scanner by the WEHI Histology Facility.

### RNA sequencing

RNA was extracted using the Quick-RNA Miniprep Kit (Zymo) as per manufacturer’s instructions. Libraries were prepared using either the Illumina TruSeq RNA Library Prep Kit v2 (Illumina) as per manufacturer’s instructions modified only to be performed using half reactions, or the Illumina®Ribo-Zero Plus rRNA depletion Kit followed by the TruSeq® Stranded Total RNA Library Prep Kit as per manufacturer’s instructions. Libraries were cleaned using AMPureXP beads (Beckman Coulter). Libraries were quantified using the Qubit DNA HS Assay Kit (Life Technologies) and then pooled and sent to the WEHI sequencing facility for sequencing with paired end reads. Initial sequencing quality control was performed with FastQC (Andrews, S. http://www.bioinformatics.babraham.ac.uk/projects/fastqc/). Adapter trimming was performed with TrimGalore! v0.6.7 (Krueger, F. http://www.bioinformatics.babraham.ac.uk/projects/trim_galore/). Reads obtained were aligned using histat2 v2.0.5 (D. Kim et al., 2015) to mm10 with SNPs between the reference genome and CAST/EiJ n-masked, created using SNPsplit (Krueger & Andrews, 2016). Reads were phased by haplotype using SNPsplit and normalised expression values in log2 reads per million (log2RPM) were determined using the Seqmonk package (www.bioinformatics.babraham.ac.uk/projects/seqmonk). Differential expression analysis was performed using EdgeR^82^ from within the Seqmonk package. t-tests for statistical analysis were performed in Microsoft Excel with Bonferroni Correction for multiple testing and Prism software (GraphPad) was used to plot data.

### ChIP sequencing

Tissue homogenisation was done by cryopulverisation of frozen samples with a pre-chilled metal mortar, pestle and hammer and the use of liquid nitrogen to maintain temperature of all equipment. Powdered tissue was resuspended in ice-cold PBS containing 1x cOmplete protease inhibitor (Roche) by pipetting and then pelleted by centrifugation at 600xg for 3 minutes at 4°C. Supernatant was discarded and tissue samples were fixed in final 1% methanol free formaldehyde (Sigma-Aldrich) in PBS for 15 minutes at room temperature with slow rotation. Samples were quenched by adding glycine (Sigma-Aldrich) to 125mM and incubating for 2 minutes at room temperature. Samples were centrifuged as previous and washed with PBS containing 1x cOmplete protease inhibitor. Samples were pelleted by centrifugation and supernatant was removed. Cells were lysed in ChIP buffer [150mM NaCl (Sigma-Aldrich), 50mM Tris-HCl (pH 7.5) (Sigma-Aldrich), 5mM EDTA (Sigma-Aldrich), 0.5% (v/v) NP-40 (Sigma-Aldrich), 1% (v/v) Triton-X (Sigma-Aldrich)] supplemented with 1X cOmplete protease inhibitor (Roche) by trituration and then incubation on ice for 10 minutes with occasional inversion. Samples were centrifuged at 12,000rpm for 1 minute at 4°C and supernatant was discarded. Samples were resuspended in MNase buffer [1x MNase buffer (NEB), 1x BSA (NEB) in nuclease free water] and pre-incubated at 37°C for 5 minutes. MNase enzyme (NEB) was added and incubated for a further 5 minutes at 37°C. 10mM EDTA (Sigma-Aldrich) was added and mixed by thorough inversion then incubated on ice for 10 minutes. Nuclei were then centrifuged at 12,000rpm for 1 minute at 4°C, supernatant was discarded, samples were resuspended in ChIP buffer and transferred to a sonication tube (Covaris). Samples were sonicated in a Covaris S220 sonicator (peak power: 75W, duty factor: 27%, cycles/burst: 200, duration: 15 seconds, temperature: 4-11°C) and then transferred back into a chilled snap cap microtube containing ChIP buffer supplemented with 1x cOmplete protease inhibitor (Roche). Samples were spun again at 12,000rpm for 1 minute at 4°C, the supernatant was retained and 10µl was set aside at 4°C to be used as whole cell extract (WCE) input control. Each immunoprecipitation was diluted to in ChIP buffer supplemented with 1x cOmplete protease inhibitor (Roche) and 10µg of antibody (anti-GFP A11122, Life Technologies). Samples were left rotating overnight at 4°C. Chromatin was cleared by centrifugation at 12,000xg for 10 minutes at 4°C and then added to 20µl of pre-washed (3x with cold ChIP buffer) Protein G DynaBeads (Thermo Fisher), which were left to incubate at 4°C rotating for 2 hours. The beads were then washed 6x with cold ChIP buffer before eluting in 1 volume of elution buffer [1% (w/v) SDS (Biorad), 0.1M NaHCO3 (Sigma-Aldrich) in Milli-Q® water] shaking at 37°C for 15 minutes. Beads were pelleted on magnets for 3 minutes, supernatant was retained. A second elution was repeated with a further 1 volume of elution buffer and was added to the first. WCE input control was diluted to 2 volumes in elution buffer and was henceforth treated the same as all samples. Samples were decrosslinked by adding 200mM NaCl and 25µg/ml RNAse A (Thermo Fisher) to each sample and incubating shaking overnight at 65°C. 100µg/ml Proteinase K (Roche) was added and the samples were incubated for 1 hour shaking at 65°C. DNA was extracted using the ChIP DNA Clean and Concentrator Kit (Zymo) as per the manufacturer’s instructions then quantified using the Qubit DNA HS Assay Kit (Life Technologies). Libraries were prepared using the Illumina TruSeq RNA Library Prep Kit v2 (Illumina) as per manufacturer’s instructions modified only to be performed using half reactions and beginning at the ‘preparing illumina adapter libraries’ step. Libraries were checked by gel electrophoresis after 15x PCR cycles and were then cleaned using AMPureXP beads (Beckman Coulter), quantified using the Qubit and pooled for sequencing. Pools were sent to the WEHI sequencing facility and sequenced using the Illumina P2 100-cycle 400M read kit, paired-end. QC of Fastq sequencing data was done using FastQC v0.11.8 (Andrews,S.http://www.bioinformatics.babraham.ac.uk/projects/fastqc/). Fastq sequencing files were processed by adapter trimming using TrimGalore! v0.6.7 (Krueger, F. http://www.bioinformatics.babraham.ac.uk/projects/trim_galore/) and Cutadapt v.3.7. Trimmed reads were mapped to the GRCm38 version of the mouse reference genome using Bowtie2 v2.3.4.1^83^ and Samtools v1.12^84^. Bam files were opened in Seqmonk version 1.48.1 and peaks were called using the Macs2^85^ from within the Seqmonk package using Smchd1 negative (no GFP) samples as controls with an FDR cut-off of 0.05.

### Home-cage analysis

Home-cage analysis was done using an automated system from Actual Analytics equipped with an RFID baseplate to track movement through individual microchips and a camera. Mice were housed 4-5 per cage, each of the regular home-cages were put into the home-cage analyser and data was recorded for 44-50 hours. Data processed on the system software, exported to excel. Prism software (GraphPad) was used for statistical analyses and to plot data.

### Grip strength testing

Tests of grip strength were performed using a grip strength meter (Bioseb) with a T-bar attachment. Mice were held by the tail, slowly lowered and allowed to grip the bar with their forelimbs and then gently pulled away until they let go. The maximum force at which they maintained their grip on the bar was recorded. This was repeated 5 times for each individual with breaks in between each testing round. The top 3 scores for each mouse were used for comparison between genotypes. Mice were weighed on the day of testing and grip strength data was normalised to body weight. Prism software (GraphPad) was used for statistical analyses and to plot data.

### Rotarod test for motor coordination

Rotarod tests were performed using the UgoBasile RotaRod for Mice (47650) on 4-5 mice (cage mates) simultaneously. Mice were placed on the rod between dividers all facing backwards and given a “ready, set, go” warning before the rod began to move, they were allowed one practise of this. Mice were given a maximum of 120 seconds at each of the following speeds: 12rpm, 20rpm, 28rpm, 36rpm and 44rpm; with a 5 minute rest between rounds. Mice were returned to their home-cage for minimum 1 hour before repetition of the test as a second trial. An HD webcam (Logitech C615) was set up to film the test and latency to fall was recorded from video footage using a stopwatch. Data was averaged across both rounds. Prism software (GraphPad) was used for statistical analyses and to plot data.

## ACKNOWLEDGEMENTS

This work was supported by National Health and Medical Research Council of Australia (NHMRC) Investigator grants [2007996 to Q.G.; 2034104 to J.M.M.; 1194345 and 2041117 to M.E.B.], a Foundation for Prader Willi Research USA grant [825553 to M.E.B.], Prader Willi Research Foundation Australia Travel Award and Research Training Program Scholarship to MI, Al and Val Rosenstrauss Fellowship from the Rebecca L. Cooper Medical Research Foundation [F20221078 to H.K.V.], Pamela and Lorenzo Galli Trust support (to M.E.B.), Brian M Davis Charitable Trust support (to M.E.B.) and support from WEHI Innovation funding. The work was additionally supported via NHMRC IRIISS and Victorian State Government Operational Infrastructure Support.

## CONFLICTS OF INTEREST

MB is a shareholder and CSO of Togglelux Therapeutics Pty Ltd, a company focused on developing novel therapeutics to treat Prader-Willi Syndrome. JM is shareholder and Director of Protein Science of Togglelux. No part of the research presented in this paper was funded or sponsored by Togglelux.

## REFERENCES

1. Bohonowych, J., Miller, J., McCandless, S.E. & Strong, T.V. The Global Prader–Willi Syndrome Registry: Development, Launch, and Early Demographics. Genes 10(2019).

2. Passone, C.B.G. et al. Prader-Willi Syndrome: What Is the General Pediatrician Supposed to Do? - a Review. Rev Paul Pediatr 36, 345–352 (2018).

3. Lionti, T., Reid, S.M., White, S.M. & Rowell, M.M. A population-based profile of 160 Australians with Prader-Willi syndrome: trends in diagnosis, birth prevalence and birth characteristics. Am J Med Genet A 167A, 371–8 (2015).

4. Alves, C. & Franco, R.R. Prader-Willi syndrome: endocrine manifestations and management. Arch Endocrinol Metab 64, 223–234 (2020).

5. Costa, R.A., Ferreira, I.R., Cintra, H.A., Gomes, L.H.F. & Guida, L.D.C. Genotype-Phenotype Relationships and Endocrine Findings in Prader-Willi Syndrome. Front Endocrinol (Lausanne) 10, 864 (2019).

6. Feighan, S.M., Hughes, M., Maunder, K., Roche, E. & Gallagher, L. A profile of mental health and behaviour in Prader-Willi syndrome. J Intellect Disabil Res 64, 158–169 (2020).

7. McCandless, S.E. Clinical report-health supervision for children with Prader-Willi syndrome. Pediatrics 127, 195–204 (2011).

8. Szabadi, S. et al. A Review of Prader–Willi Syndrome. Endocrines 3, 329–348 (2022).

9. Miller, J.L. et al. Nutritional phases in Prader-Willi syndrome. Am J Med Genet 155A, 1040–9 (2011).

10. Chen, C., Visootsak, J., Dills, S. & Graham, J.M. Prader-Willi Syndrome: An Update and Review for the Primary Pediatrician. Clinical Pediatrics 46, 580–591 (2007).

11. Horsthemke, B. & Wagstaff, J. Mechanisms of imprinting of the Prader-Willi/Angelman region. Am J Med Genet A 146A, 2041–52 (2008).

12. Butler, M.G. et al. Molecular genetic classification in Prader-Willi syndrome: a multisite cohort study. J Med Genet 56, 149–153 (2019).

13. Beygo, J. et al. Common genetic variation in the Angelman syndrome imprinting centre affects the imprinting of chromosome 15. Eur J Hum Genet (2020).

14. Reis, A. et al. Imprinting Mutations Suggested by Abnormal DNA Methylation Paierns in Familial Angelman and Prader-Willi Syndromes. Am J Hum Genet 54, 741–747 (1994).

15. Cruvinel, E. et al. Reactivation of maternal SNORD116 cluster via SETDB1 knockdown in Prader-Willi syndrome iPSCs. Human Molecular Genetics 23, 4674–4685 (2014).

16. S Saitoh et al. Minimal definition of the imprinting center and fixation of chromosome 15q11-q13 epigenotype by imprinting mutations. Proc Natl Acad Sci U S A 93, 7811–7815 (1996).

17. Ariyanfar, S. & Good, D.J. Analysis of SNHG14: A Long Non-Coding RNA Hosting SNORD116, Whose Loss Contributes to Prader-Willi Syndrome Etiology. Genes (Basel) 14(2023).

18. Cavaillé, J. et al. Identification of brain-specific and imprinted small nucleolar RNA genes exhibiting an unusual genomic organization. PNAS 97, 14311–14316 (2000).

19. Hsiao, J.S. et al. A bipartite boundary element restricts UBE3A imprinting to mature neurons. Proc Natl Acad Sci U S A 116, 2181–2186 (2019).

20. Meng, L., Person, R.E. & Beaudet, A.L. Ube3a-ATS is an atypical RNA polymerase II transcript that represses the paternal expression of Ube3a. Hum Mol Genet 21, 3001–12 (2012).

21. LaSalle, J.M., Reiter, L.T. & Chamberlain, S.J. Epigenetic regulation of UBE3A and roles in human neurodevelopmental disorders. Epigenomics 7, 1213–1228 (2015).

22. Kummerfeld, D.M. et al. A Comprehensive Review of Genetically Engineered Mouse Models for Prader-Willi Syndrome Research. Int J Mol Sci 22(2021).

23. Saitoh, S. & Wada, T. Parent-of-Origin Specific Histone Acetylation and Reactivation of a Key Imprinted Gene Locus in Prader-Willi Syndrome. American Journal of Human Genetics 66, 1958–1962 (2000).

24. Kim, Y. et al. Targeting the histone methyltransferase G9a activates imprinted genes and improves survival of a mouse model of Prader–Willi syndrome. Nature Medicine 23, 213–224 (2017).

25. Langouët, M. et al. Specific ZNF274 binding interference at SNORD116 activates the maternal transcripts in Prader-Willi syndrome neurons. Hum Mol Genet 29, 3285–3295 (2020).

26. Langouët, M. et al. Zinc finger protein 274 regulates imprinted expression of transcripts in Prader-Willi syndrome neurons. Human Molecular Genetics 27, 505–515 (2018).

27. Wang, S. et al. Newly developed oral bioavailable EHMT2 inhibitor as a potential epigenetic therapy for Prader-Willi syndrome. Mol Ther 32, 2662–2675 (2024).

28. Wu, H. et al. Small molecule inhibitors of G9a reactivate the maternal PWS genes in Prader-Willi-Syndrome patient derived neural stem cells and differentiated neurons. bioRxiv May(2019).

29. Rohm, D. et al. Activation of the imprinted Prader-Willi syndrome locus by CRISPR-based epigenome editing. Cell Genom 5, 100770 (2025).

30. Griazeva, E.D. et al. Current Approaches to Epigenetic Therapy. Epigenomes 7(2023).

31. Dindot, S.V. et al. An ASO therapy for Angelman syndrome that targets an evolutionarily conserved region at the start of the UBE3A-AS transcript. Science Translational Medicine 15(2023).

32. Meng, L. et al. Towards a therapy for Angelman syndrome by targeting a long non-coding RNA. Nature 518, 409–12 (2015).

33. Milazzo, C., et al. Antisense oligonucleotide treatment rescues UBE3A expression and multiple phenotypes of an Angelman syndrome mouse model. JCI Insight 6(2021).

34. Hipp, J.F. et al. The UBE3A-ATS antisense oligonucleotide rugonersen in children with Angelman syndrome: a phase 1 trial. Nat Med 31, 2936–2945 (2025).

35. Sahoo, T. et al. Prader-Willi phenotype caused by paternal deficiency for the HBII-85 C/D box small nucleolar RNA cluster. Nat Genet 40, 719–21 (2008).

36. de Smith, A.J. et al. A deletion of the HBII-85 class of small nucleolar RNAs (snoRNAs) is associated with hyperphagia, obesity and hypogonadism. Hum Mol Genet 18, 3257–65 (2009).

37. Duker, A.L. et al. Paternally inherited microdeletion at 15q11.2 confirms a significant role for the SNORD116 C/D box snoRNA cluster in Prader-Willi syndrome. Eur J Hum Genet 18, 1196–201 (2010).

38. Grootjen, L.N. et al. Prenatal and Neonatal Characteristics of Children with Prader-Willi Syndrome. J Clin Med 11(2022).

39. Fontana, P. et al. SNORD116 deletions cause Prader-Willi syndrome with a mild phenotype and macrocephaly. Clin Genet 92, 440–443 (2017).

40. Tan, Q. et al. Prader-Willi-Like Phenotype Caused by an Atypical 15q11.2 Microdeletion. Genes (Basel) 11(2020).

41. Bieth, E. et al. Highly restricted deletion of the SNORD116 region is implicated in Prader-Willi Syndrome. Eur J Hum Genet 23, 252–5 (2015).

42. Schaaf, C.P. et al. Truncating mutations of MAGEL2 cause Prader-Willi phenotypes and autism. Nat Genet 45, 1405–8 (2013).

43. Schubert, T. & Schaaf, C.P. MAGEL2 (patho-)physiology and Schaaf-Yang syndrome. Dev Med Child Neurol 67, 35–48 (2024).

44. McCarthy, J.M. et al. Hormonal, metabolic and skeletal phenotype of Schaaf-Yang syndrome: a comparison to Prader-Willi syndrome. J Med Genet 55, 307–315 (2018).

45. McCarthy, J. et al. Schaaf-Yang syndrome overview: Report of 78 individuals. Am J Med Genet A 176, 2564–2574 (2018).

46. Abreu, A.P. et al. Central precocious puberty caused by mutations in the imprinted gene MKRN3. N Engl J Med 368, 2467–75 (2013).

47. Brideau, N.J. et al. Independent Mechanisms Target SMCHD1 to Trimethylated Histone H3 Lysine 9-Modified Chromatin and the Inactive X Chromosome. Mol Cell Biol 35, 4053–68 (2015).

48. Chen, K. et al. Crystal structure of the hinge domain of Smchd1 reveals its dimerization mode and nucleic acid–binding residues. Science Signaling 13(2020).

49. Chen, K. et al. The epigenetic regulator Smchd1 contains a functional GHKL-type ATPase domain. Biochem J 473, 1733–44 (2016).

50. Chen, K., Czabotar, P.E., Blewitt, M.E. & Murphy, J.M. The hinge domain of the epigenetic repressor Smchd1 adopts an unconventional homodimeric configuration. Biochem J 473, 733–42 (2016).

51. Chen, K. et al. Genome-wide binding and mechanistic analyses of Smchd1-mediated epigenetic regulation. Proc Natl Acad Sci U S A 112, E3535–44 (2015).

52. Gurzau, A.D. et al. SMCHD1’s ubiquitin-like domain is required for N-terminal dimerization and chromatin localization. Biochem J (2021).

53. Gurzau, A.D., Blewitt, M.E., Czabotar, P.E., Murphy, J.M. & Birkinshaw, R.W. Relating SMCHD1 structure to its function in epigenetic silencing. Biochem Soc Trans 48, 1751–1763 (2020).

54. Pedersen, L.C., Inoue, K., Kim, S., Perera, L. & Shaw, N.D. A ubiquitin-like domain is required for stabilizing the N-terminal ATPase module of human SMCHD1. Commun Biol 2, 255 (2019).

55. Benetti, N., et al. Maternal SMCHD1 regulates Hox gene expression and patterning in the mouse embryo. Nat Commun 13, 4295 (2022).

56. Gendrel, A.V. et al. Epigenetic functions of smchd1 repress gene clusters on the inactive X chromosome and on autosomes. Mol Cell Biol 33, 3150–65 (2013).

57. Jansz, N. et al. Smchd1 Targeting to the Inactive X Is Dependent on the Xist-HnrnpK-PRC1 Pathway. Cell Rep 25, 1912–1923 e9 (2018).

58. Jansz, N. et al. Smchd1 regulates long-range chromatin interactions on the inactive X chromosome and at Hox clusters. Nat Struct Mol Biol 25, 766–777 (2018).

59. Mould, A., et al. Smchd1 regulates a subset of autosomal genes subject to monoallelic expression in addition to being critical for X inactivation. Epigenetics Chromatin. 6(2013).

60. Wanigasuriya, I. et al. Smchd1 is a maternal effect gene required for genomic imprinting. Elife 9(2020).

61. Kinkel, S.A. et al. Epigenetic modifier SMCHD1 maintains a normal pool of long-term hematopoietic stem cells. iScience 25, 104684 (2022).

62. Leong, H.S. et al. Epigenetic regulator Smchd1 functions as a tumor suppressor. Cancer Res 73, 1591–9 (2013).

63. Tronche, F.o., et al. Disruption of the glucocorticoid receptor gene in the nervous system results in reduced anxiety. Nature Genetics 23, 99–103 (1999).

64. de Greef, J.C. et al. Smchd1 haploinsufficiency exacerbates the phenotype of a transgenic FSHD1 mouse model. Hum Mol Genet 27, 716–731 (2018).

65. Blewii, M.E. et al. SmcHD1, containing a structural-maintenance-of-chromosomes hinge domain, has a critical role in X inactivation. Nat Genet 40, 663–9 (2008).

66. Kozlov, S.V. et al. The imprinted gene Magel2 regulates normal circadian output. Nature Genetics 39, 1266–1272 (2007).

67. Ventura, A. et al. Restoration of p53 function leads to tumour regression in vivo. Nature 445, 661–5 (2007).

68. Luo, L. et al. Optimizing Nervous System-Specific Gene Targeting with Cre Driver Lines: Prevalence of Germline Recombination and Influencing Factors. Neuron 106, 37–65 e5 (2020).

69. Zhang, J. et al. Germ-line recombination activity of the widely used hGFAP-Cre and nestin-Cre transgenes. PLoS One 8, e82818 (2013).

70. Tapia Del Fierro, A., et al. SMCHD1 has separable roles in chromatin architecture and gene silencing that could be targeted in disease. Nat Commun 14, 5466 (2023).

71. Fountain, M.D., Tao, H., Chen, C.A., Yin, J. & Schaaf, C.P. Magel2 knockout mice manifest altered social phenotypes and a deficit in preference for social novelty. *Genes*, Brain and Behaviour 16, 592–600 (2017).

72. Kamaludin, A. et al. Muscle dysfunc9on caused by loss of Magel2 in a mouse model of Prader-Willi and Schaaf-Yang syndromes. Human Molecular Genetics 25, 3798–3809 (2016).

73. Mercer, R. & Wevrick, R. Loss of Magel2, a Candidate Gene for Features of Prader-Willi Syndrome, Impairs Reproductive Function in Mice. PloS ONE 4, 1–9 (2009).

74. Bischof, J., Stewart, C.L. & Wevrick, R. Inactivation of the mouse Magel2 gene results in growth abnormalities similar to Prader-Willi syndrome. Human Molecular Genetics 16, 2713–2719 (2007).

75. Wolff, R., et al. The Preclinical Animal Network (PCAN): Integrative high-throughput phenotyping of standardized mouse models for Prader-Willi syndrome. bioRxiv (2025).

76. Bains, R.S. et al. Analysis of Individual Mouse Activity in Group Housed Animals of Different Inbred Strains using a Novel Automated Home Cage Analysis System. Front Behav Neurosci 10, 106 (2016).

77. Cataldi, M. et al. Sleep disorders in Prader-Willi syndrome, evidence from animal models and humans. Sleep Med Rev 57, 101432 (2021).

78. Pavone, M. et al. Sleep disordered breathing in patients with Prader-Willi syndrome: A multicenter study. Pediatr Pulmonol 50, 1354–9 (2015).

79. Yang, T. et al. A mouse model for Prader-Willi syndrome imprinting-centre mutations. Nature Genetics 19, 25–31 (1998).

80. Chapman, V. & Ruddell, F. Glutamate Oxaloacetate Transaminase (GOT) genetics in the mouse: polymorphism of GOT-1. Genetics 70, 299–305 (1971).

81. Petkov, P.M. et al. Development of a SNP genotyping panel for genetic monitoring of the laboratory mouse. Genomics 83, 902–11 (2004).

82. Robinson, M.D., McCarthy, D.J. & Smyth, G.K. edgeR: a Bioconductor package for differential expression analysis of digital gene expression data. Bioinformatics 26, 139–40 (2010).

83. Langmead, B. & Salzberg, S.L. Fast gapped-read alignment with Bowtie 2. Nat Methods 9, 357–9 (2012).

84. Li, H. et al. The Sequence Alignment/Map format and SAMtools. Bioinformatics 25, 2078–9 (2009).

85. Zhang, Y. et al. Model-based analysis of ChIP-Seq (MACS). Genome Biol 9, R137 (2008).

