## Supplementary Figures for "SMCHD1 is a novel target for gene-activation therapy to treat Prader-Willi Syndrome"

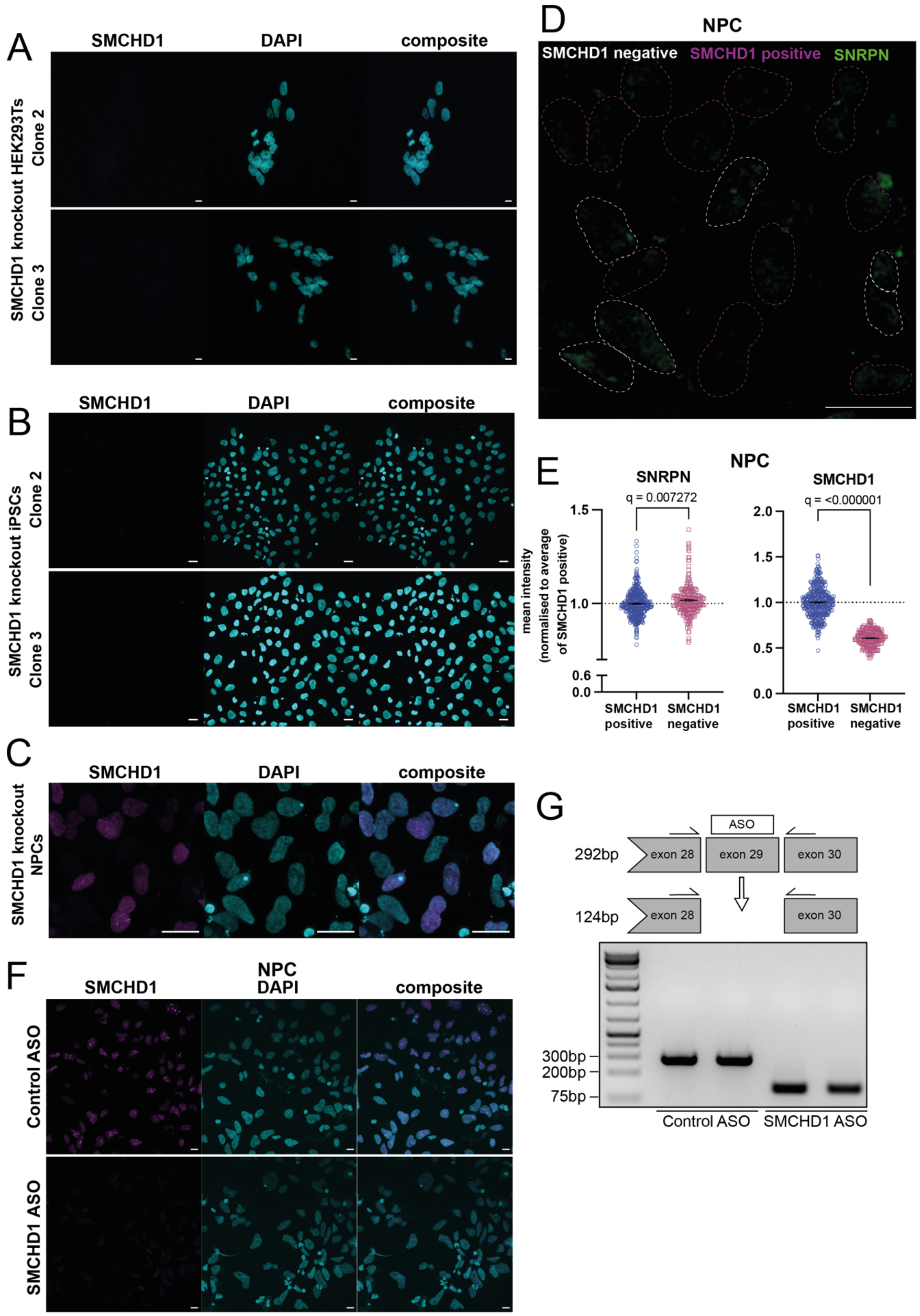

A
SMCHD1
DAPI
composite
SMCHD1 knockout HEK293Ts
Clone 2
Clone 3
B
SMCHD1
DAPI
composite
SMCHD1 knockout iPSCs
Clone 2
Clone 3
C
SMCHD1
DAPI
composite
SMCHD1 knockout NPCs
D
NPC
SMCHD1 negative
SMCHD1 positive
SNRPN
E
NPC
SNRPN
q = 0.007272
SMCHD1
q = <0.000001
mean intensity (normalised to average of SMCHD1 positive)
1.5
1.0
0.6
0.0
2.0
1.5
1.0
0.5
0.0
SMCHD1 positive
SMCHD1 negative
SMCHD1 positive
SMCHD1 negative
F
SMCHD1
NPC DAPI
composite
Control ASO
SMCHD1 ASO
G
ASO
292bp
exon 28
exon 29
exon 30
124bp
exon 28
exon 30
300bp
200bp
75bp
Control ASO
SMCHD1 ASO

**Supplementary Figure 1: SMCHD1 knockout in clonal human cell lines and in patient-derived neural cells.** Representative confocal images showing SMCHD1 protein by immunofluorescence, SMCHD1 in magenta, DAPI in cyan, scale bar 20$\mu$m. **A)** In two further HEK293T clonal lines **B)** In two further clonal PWS mUPD iPSC lines **C/F)** In PWS neural progenitors (**C**) post nucleofection of CRISPR ribonucloprotein complexes or (**F**) post treatment with either an ASO targeting SMCHD1 or a control ASO for 72 hours. **D)** Representative confocal image of SNRPN protein (green) by immunofluorescence in NPCs derived from PWS large deletion iPSCs nucleofected with CRISPR ribonucleoproteins, individual nuclei outlined in white (SMCHD1 knockout) or magenta (SMCHD1 wildtype), scale bar 20$\mu$m. **E)** Plots of mean SNRPN (left) and SMCHD1 (right) fluorescence intensity for each scored nucleus normalised to the average intensity in SMCHD1 wild-type cells from the same image n=410 wild-type nuclei, n=229 knockout nuclei from two separate differentiations and nucleofections, t-tests with Benjamini and Hochberg corrections for multiple testing, $q < 0.01$ for significance. **G)** RT-PCR products run on 2% agarose gel representative of exon skipping after 72h ASO treatment with schematic of expected band sizes for successful SMCHD1 exon skipping.

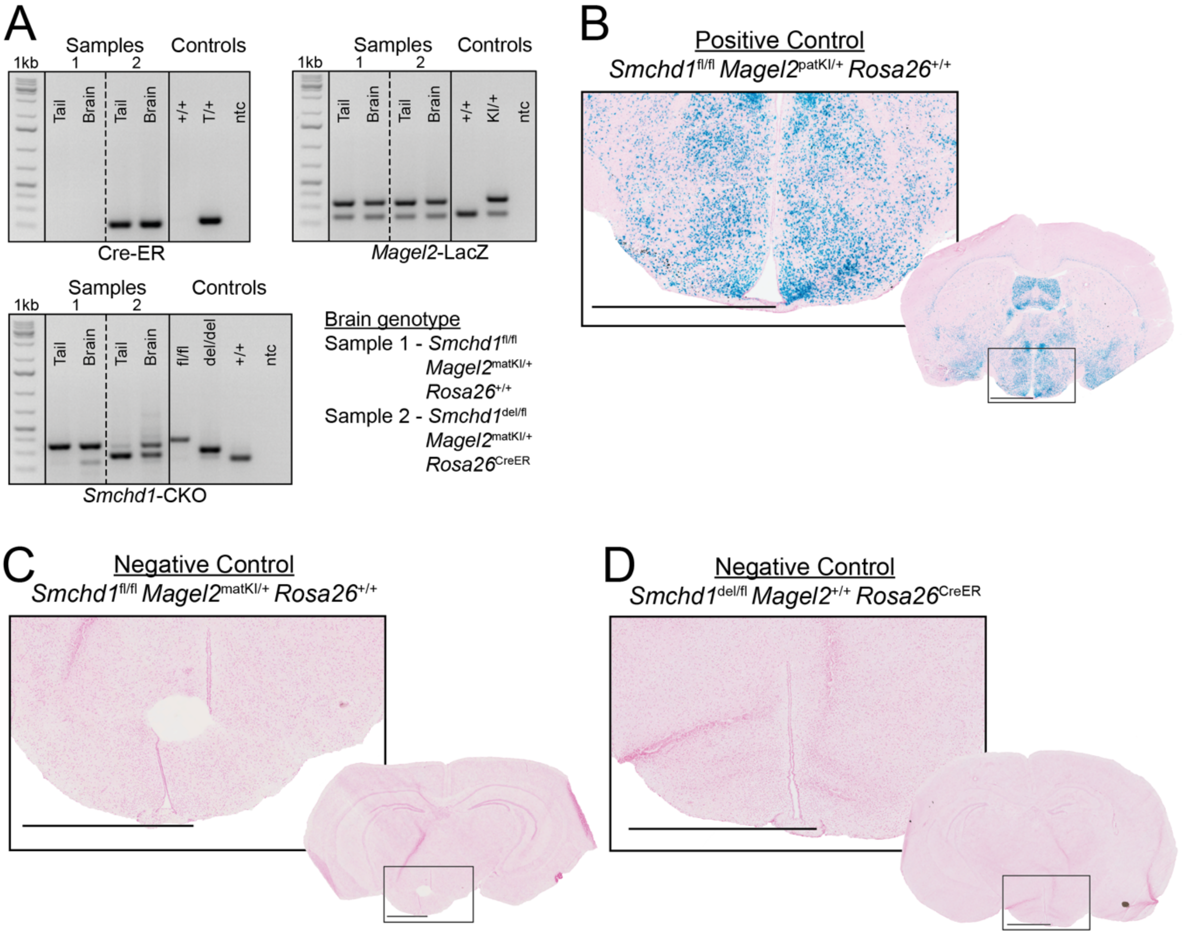


**Supplementary Figure 2: Partial deletion of *Smchd1* in the brain by tamoxifen-inducible CreERT2 with expected *Magel2* expression in all *Magel2* control brain cryosections with wild-type Smchd1. A)** Representative set of genotyping PCR products run on 1.8% w/v agarose gel using DNA purified from tail and brain samples collected at time of harvest, all PCR products run on the same gel, dotted line separates adjacent samples, solid line indicates cropped out samples. **B-D)** Representative images of brain cryosections stained with X-gal showing proxy for *Magel2* expression (blue) and counterstained with nuclear fast red (pink), zoomed images correspond to their overlayed whole brain slice, scale bars 1500$\mu$m. Note vertical brain sections are not matched exactly by slice.

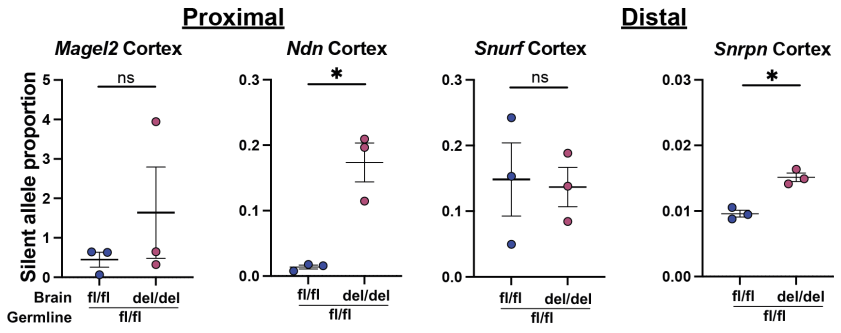


**Supplementary Figure 3: *Nestin*-Cre deletion of *Smchd1 in vivo* leads to activation of both proximal and distal maternal PWS cluster genes in the cortex.** Graphs showing proportion of PWS gene expression from the imprinted maternal allele as a proportion of expression from the paternal allele by RNA sequencing in female mice, t-test with Bonferroni correction for multiple testing, ns = not significant * = FDR <0.05.

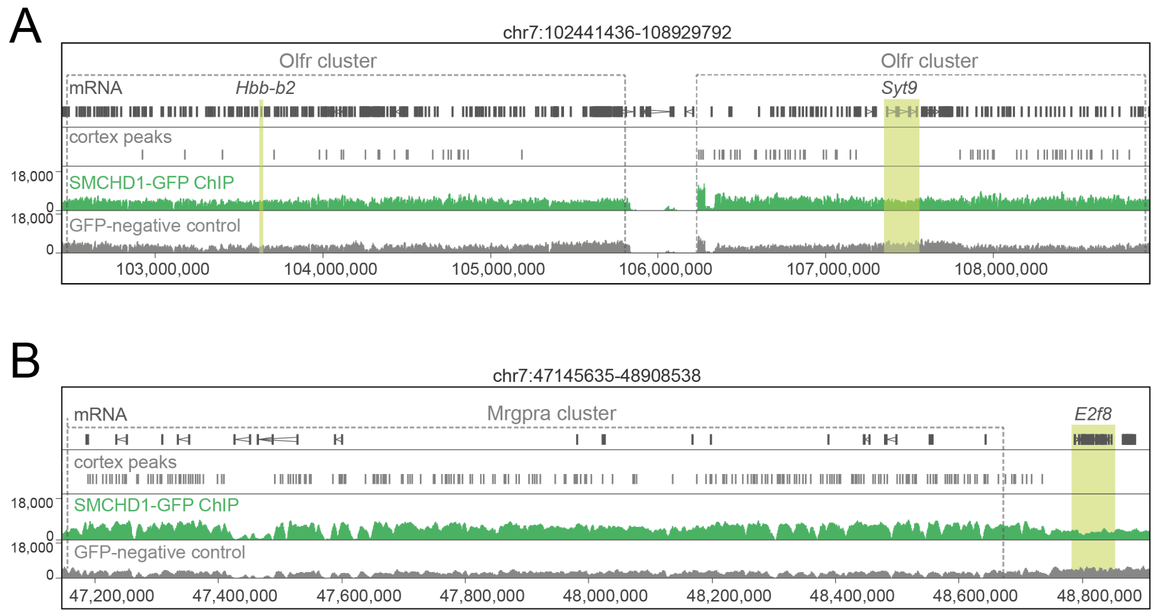


**Supplementary Figure 4: SMCHD1 binding at DE genes in mouse cortex tissue.** Genome browser ChIP-seq tracks for SMCHD1-GFP ChIP-seq showing peaks at (**A**) olfactory receptor gene clusters and (**B**) the *Mgrpra* gene cluster on chromosome 7 with *Hbb-b2*, *Syt9* and *E2f8* highlighted as differentially expressed between *Smchd1*$^{del/del}$ and *Smchd1*$^{fl/fl}$ cortex samples by RNA-seq, n=2 for each sample.

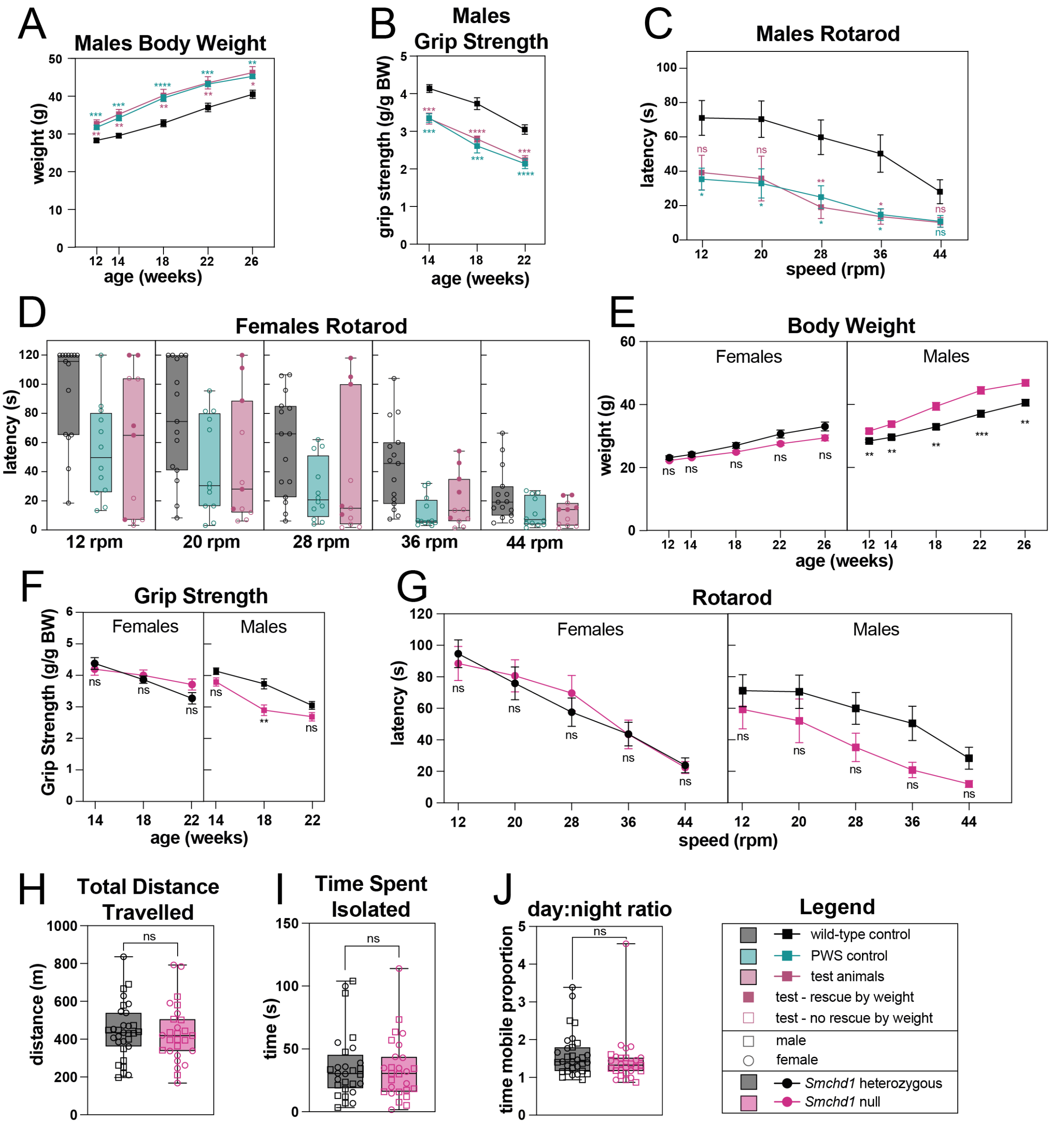


**Supplementary Figure 5: *Nestin*-Cre deletion of *Smchd1* has minimal effects on male mice *Magel2*$^{\text{patKI/+}}$ mice and on *Magel2* replete mice of both sexes.** n=11-16 per genotype across two cohorts. **A)** Total body weight in male mice. **B)** grip strength in male mice normalised to body weight. **C)** Average latency to fall from two rounds of testing for motor coordination on a rotarod in male mice. **D)** Average latency to fall plotted per individual for female mice from two rounds of rotarod testing for motor coordination. **E)** Total body weight in female and male mice. **F)** Grip strength measurements normalised to body weight in female and male mice. **G)** Average latency to fall from two rounds of testing for motor coordination on a rotarod in female and male mice. **H-J)** Data from home-cage observation periods of >44h for both male and female mice combined across two cohorts showing (**H**)Total distance travelled (**I**) Average time spent isolated from all other cage-mates per hour from two combined 10h active periods, each cohort normalised to average of wildtype for statistical analysis. (**J**) Plot of the ratio of time spent mobile in the active versus inactive periods.

Genotypes in the brain are wild-type control = $Smchd1^{del/fl}Magel2^{+/+}$, PWS control = $Smchd1^{del/fl}Magel2^{patKI/+}$, test animals = $Smchd1^{del/del}Magel2^{patKI/+}$, error bars = ±SEM, **(A-G)** two-way ANOVA Tukey's multiple comparison or **(H-J)** Welch's t-test adjusted p-values ns = not significant, ** = $p < 0.01$, *** = $p < 0.001$.
