## Supplementary Table 1 for "SMCHD1 is a novel target for gene-activation therapy to treat Prader-Willi Syndrome"

| **Guide** | **Coordinates** | **Protein generated** |
| --- | --- | --- |
| 1-1 | 2,656,147 – 2,656,169 | **wildtype:**  EEDGGGVGHRTVYLFD  **gRNA(15%):**  EEDGGGVGHSVLV*STOP* |
| 3-2 | 2,666,868 – 2,667,067 | **wildtype:**  LPHYDTLVKSG  **gRNA(22%):**  LPHYDTG*STOP* |
