## Supplementary Table 2 for "SMCHD1 is a novel target for gene-activation therapy to treat Prader-Willi Syndrome"

| **Name** | **Patient** | **Genetics** | **Sex** | **Reference iPSC** | **Source** |
| --- | --- | --- | --- | --- | --- |
| PWS 1.7 LD | PWS | Large deletion BP2-BP3 | XX | S. J. Chamberlain  *et al*., 2010 | UCONN stem cell core |
| PWS 1.2 UPD | PWS | matUPD | XX | Langouët *et al*.,  2018 | UCONN stem cell core |
| YK27 | Control | No mutation | XX | Martins-Taylor  *et al*., 2011 | UCONN stem cell core |
