## Supplementary Table 14 for "SMCHD1 is a novel target for gene-activation therapy to treat Prader-Willi Syndrome"

| **Name** | **Forward sequence** | **Reverse sequence** | **Use** |
| --- | --- | --- | --- |
| *SMCHD1* ASO | GUCCAGAAAUUAGUUGCACUC (Lemmers *et al.*, 2012) | | ASO |
| Control ASO  (targeting Dystrophin) | GGCUGCUUUGCCCUC (Lemmers *et al.*, 2012) | | ASO |
| SMCHD1 skipping | CTGGGGTTGGACTTGATAGC | GGTGCTGGATTATCCCACTG | RT-PCR |
| SMCHD1 guide 1-1 | GTGACCTATGAACTCAGGAGTCtcgcgtacctgacacacaca | CTGAGACTTGCACATCGCAGCcgctgtcttttctccttttc | MiSeq |
| SMCHD1 guide 3-2 | GTGACCTATGAACTCAGGAGTCtcttgattacagatgaaact | CTGAGACTTGCACATCGCAGCaaggcaggtaataatttatc | MiSeq |
| SMCHD1 crRNA | CAAACAAGTACACCGTCCTG | | CRISPR RNP |
| hSMCHD1.1 shRNA | TGCTGTTGACAGTGAGCGCCAAGTTGAAGAAGCAAGATTATAGTGAAGCCACAGATGTATAATCTTGCTTCTTCAACTTGATGCCTACTGCCTCGGA | | shRNA |
| hSMCHD1.2 shRNA | TGCTGTTGACAGTGAGCGCCCGGTACAACATGTTAAAATATAGTGAAGCCACAGATGTATATTTTAACATGTTGTACCGGTTGCCTACTGCCTCGGA | | shRNA |
| Non-silencing shRNA | TGCTGTTGACAGTGAGCGATCTCGCTTGGGCGAGAGTAAGTAGTGAAGCCACAGATGTACTTACTCTCGCCCAAGCGAGAGTGCCTACTGCCTCGGA | | shRNA |
| Cre | CTGACCGTACACCAAAATTTGCCTG | GATAATCGCGAACATCTTCAGGTTC | Genotyping |
| Magel2 MUT | ATGGCTCCATCAGGAGAAC | GGGATAGGTCACGTTGGTGT | Genotyping |
| Magel2 WT |  | GATGGAAAGACCCTTGAGGT | Genotyping |
| Smchd1 DEL | GTAGCGCCAAGTGCCCAG | GGACAGCCAAAGTGACACAG | Genotyping |
| Smchd1 WT + FL | TCAGGTGGTCTCGAGCCC |  | Genotyping |
| MAGEL2 | GAGGTCGCAAGTGTCTCTCC | ACTTTCACCATCTCCGAGCG | RT-qPCR |
| SNRPN | CTTCTGCCCAGCTTGCAT | TGAAGATTCGGCCATCTTGC | RT-qPCR |
| GUS | CTCATTTGGAATTTTGCCGATT | CCGAGTGAAGATCCCCTTTTTA | RT-qPCR |
